# The vacuolar iron transporter mediates iron detoxification in *Toxoplasma gondii*

**DOI:** 10.1101/2021.09.08.458725

**Authors:** Dana Aghabi, Megan Sloan, Zhicheng Dou, Alfredo J. Guerra, Clare R. Harding

**Affiliations:** Wellcome Centre for Integrative Parasitology, Institute of Infection, Immunity and Inflammation, University of Glasgow, UK; Department of Biological Sciences, Clemson University, Clemson, South Carolina, USA; Department of Microbiology and Immunology, University of Michigan, Ann Arbor, Michigan, USA

## Abstract

Iron is essential to living cells, acting as a cofactor in a number of essential enzymes in metabolism; however, iron requires proper storage or it can be dangerous to the cell. In both yeast and plants, iron is stored in a vacuole through the action of a vacuolar iron transporter (VIT). This transporter is conserved in the apicomplexan family of obligate intracellular parasites, including in *Toxoplasma gondii*, a pathogen of medical and veterinary importance. Here, we assess the role of VIT in *T. gondii*. We show that deletion of VIT causes a slight growth defect *in vitro*, however leads to hypersensitivity in the presence of excess iron, confirming its essential role in iron detoxification in the parasite. In the absence of VIT, parasites contain less iron and are at a growth disadvantage when moving into an iron-depleted environment. We show parasite VIT expression is regulated by environmental iron levels at both the transcript and protein level, and by altering the distribution of VIT within the cell. In the absence of VIT, we find that the *T. gondii* responds by altering expression of genes with a role in iron metabolism and by increasing the activity of the antioxidant protein catalase. We also show that iron detoxification has an important role both in parasite survival within macrophages and in pathogenesis in a mouse model. Together, by demonstrating a critical role for VIT during iron detoxification in *T. gondii*, we reveal the importance of iron storage in the parasite and provide the first insight into the machinery involved.

## Introduction

Iron plays a central role in proteins essential for respiration in almost all living cells. However, the ease with which it gains and loses electrons makes it potentially dangerous in the cellular environment (Galaris et al., 2019; Imlay et al., 1988). Free iron reacts with H_2_O_2_, produced by aerobic respiration, via the Fenton reaction to form highly reactive hydroxyl and hydroperoxyl radicals (HO· and HOO·) which can cause extensive damage to the cell. To avoid this fate, mammalian cells safely store the majority of iron in the cytosol within protein cages of ferritin (Arosio et al., 2015). Ferritin has been highly conserved through evolution, with homologs found in plants and bacteria (Arosio et al., 2017). However, outside of the Metazoa, organisms including plants, yeast and protists instead transport iron into membrane-bound vacuoles through the action of an H^+^-dependent antiporter, named CCC1 in yeast or the vacuolar iron transporter (VIT) in plants (Li et al., 2001; Sorribes-Dauden et al., 2020). The absence of these transporters in animal cells and their importance across pathogenic fungi and protists, makes these transporters attractive targets for drug discovery (Sorribes-Dauden et al., 2020).

In common with most eukaryotes, elemental analysis found that around 5% of the atoms of *T. gondii* atoms are iron (Al-sandaqchi et al., 2018). This iron is required for a number of essential cellular processes, including heme biosynthesis (Bergmann et al., 2020; Kloehn et al., 2020), iron sulphur (Fe-S) cluster biogenesis (Aw et al., 2020), and as a cofactor in several important proteins such as catalase (Kwok et al., 2004; Odberg-Ferragut et al., 2000), prolyl hydroxylases (Florimond et al., 2019) and the deoxyribonucleotide synthesis enzyme ribonucleotide reductase. Iron acquisition and use have been studied in various pathogenic parasites such as *Trichomonas* (Sehgal et al., 2012) and in some detail in kinetoplastids such as *Leishmania* (Zaidi et al., 2017) and *Trypanosoma* (Carbajo et al., 2021). However, iron utilisation remains understudied in the apicomplexan parasites. Although the chloroquine resistance transporter (CRT) (Warring et al., 2014) was proposed to transport iron (Bakouh et al., 2017), it appears likely that its main role in *Plasmodium* is in transporting small peptides (Shafik et al., 2020). Instead, a homologue of the plant iron transporter VIT1 was described and shown to be a functional iron transporter that complemented a Δ*ccc1* yeast strain (Labarbuta et al., 2017; Sharma et al., 2021; Slavic et al., 2016). Deletion of VIT in *Plasmodium* demonstrated that the gene was essential for replication in both blood and liver stage parasites. Further, absence of VIT rendered *Plasmodium* more susceptible to iron overload and more resistant to iron removal (Slavic et al., 2016). However, *Plasmodium* spp. undergo their asexual lifecycles in the unusual environment of the red blood cell, with very high levels of iron in the form of haemoglobin. In contrast, the ubiquitous pathogen *T. gondii* is a generalist, able to infect almost any warm blooded species and any nucleated cell, exposing it to a range in iron availability. We were interested in how *T. gondii* stores iron, and the effects disrupting iron storage has on the parasite.

Based on homology to the *Plasmodium* and plant vacuolar iron transporters, we identified VIT in *T. gondii* and investigated the role of VIT and iron storage in the parasite. We find that although VIT is not essential for replication, expression of VIT confers a survival advantage during *in vitro* culture. We show that deletion of VIT makes the parasites significantly more sensitive to exogenous iron, and that this effect is mediated through an increase in reactive oxygen species (ROS) upon iron treatment. VIT has a highly dynamic localisation, changing throughout the cell cycle. We see transient colocalisation with a marker of the vacuolar associated compartment (VAC), suggesting this as the location of stored iron. Further, VIT expression is regulated by the presence or absence of exogenous iron in *T. gondii*. We also find that parasites lacking VIT are more susceptible to killing by activated macrophages, and this is likely a key factor in their inability to cause pathogenesis in mice.

## Results

### Deletion of VIT leads to a moderate growth defect

The *T. gondii* VIT was previously identified through homology to plant and *Plasmodium* transporters (Slavic et al., 2016). To confirm the conservation of key residues, we aligned the protein sequence of *T. gondii* VIT (TGGT1_266800) with sequences from *Plasmodium*, yeast (CCC1) and plants (*Arabidopsis thaliana*, where VIT1 was first identified and *Eucalyptus grandis*, as the crystal structure is available (Kato et al., 2019)) (**Fig. 1A**). The *T. gondii* VIT had 33 % identity with AtVIT1, including conservation of residues D43 and M80, recently shown to be essential for iron binding (Kato et al., 2019). To investigate the function of VIT in *T. gondii*, the coding region was deleted using two small guide RNAs targeting the 5’ and 3’ of the open reading frame (see **Table 1**) and replaced with a cassette encoding mNeonGreen under the control of the SAG1 promotor (**Fig. 1B**) in an ΔKu80 parental cell line. Integration of the cassette and loss of the native ORF was confirmed by PCR (**Fig. 1B**). ΔVIT parasites were viable in culture as they were able to form plaques, or cleared areas on a monolayer of host cells, at 6 days post infection (**Fig. 1C**). A genome wide essentiality screen predicted that loss of VIT had a mild effect on parasite growth (phenotype score -1.22) (Sidik et al., 2016). To examine the growth of the parasites in more detail, we quantified the number of parasites within a parasitophorous vacuole at 24 h post infection (**Fig. 1D**). The ΔVIT line had significantly (at the 2 cell, *p* = 0.0005 and the 8 cell, *p* = 0.0037, multiple *t* tests corrected by Holm-Sidak) fewer parasites per vacuole, demonstrating that VIT has a role in parasite replication in normal growth conditions. We also examined extracellular survival. We mechanically released parasites from host cells and incubated extracellularly at 37 °C for the indicated time before allowing them to invade fresh cells and quantifying the number of plaques formed. We found no difference in extracellular survival between the parental and ΔVIT parasites (*p* > 0.05, multiple *t* tests corrected by Holm-Sidak) (**Fig. 1E**), demonstrating that VIT does not have a role in parasite survival outside of the host cell.

**Figure 1.**
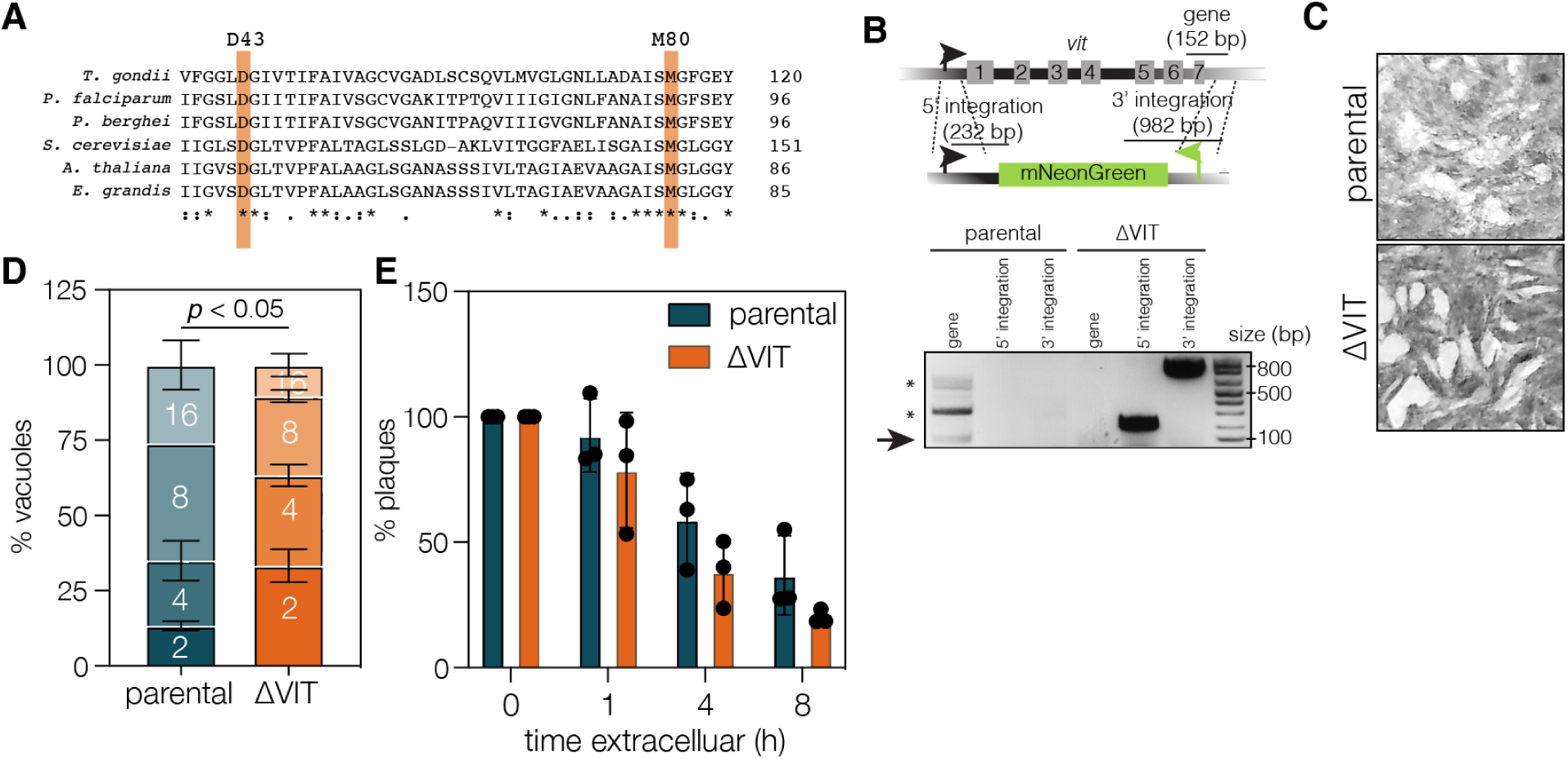
Construction and analysis of a ΔVIT parasite line. **A**. Alignment of VIT1 using Clustal Omega from *T. gondii* (TGGT1_266800), *P. falciparum* (PF3D7_1223700), *P. berghei* (PBANKA_1438600), *S. cerevisiae* (CCC1), *A. thaliana* (AtVIT1) *and E. grandis* (EgVIT1). Identity (*) and similarity (., :) indicated. Key residues for iron binding highlighted in orange. **B**. Schematic indicating the method used to replace the endogenous *vit* gene with the mNeonGreen cassette. PCR reactions confirming the replacement of the endogenous gene, expected sizes indicated on schematic. * represents secondary bands, possibly unspecific. **B**. Plaque assay demonstrating that the ΔVIT line is viable. Representative of two independent experiments. **C**. Quantification of number of parasites/vacuole at 24 h post infection. Results are the mean of three independent experiments, ± SD. *p* value from multiple *t* tests, corrected by Holm-Sidak. **D**. Quantification of extracellular survival. There was no change in the percentage of plaques formed after incubation at various times extracellularly (normalised to 0 h) between the parental and the ΔVIT parasite line. Bars are the mean of three independent experiments, ± SD.

**Table 1.**
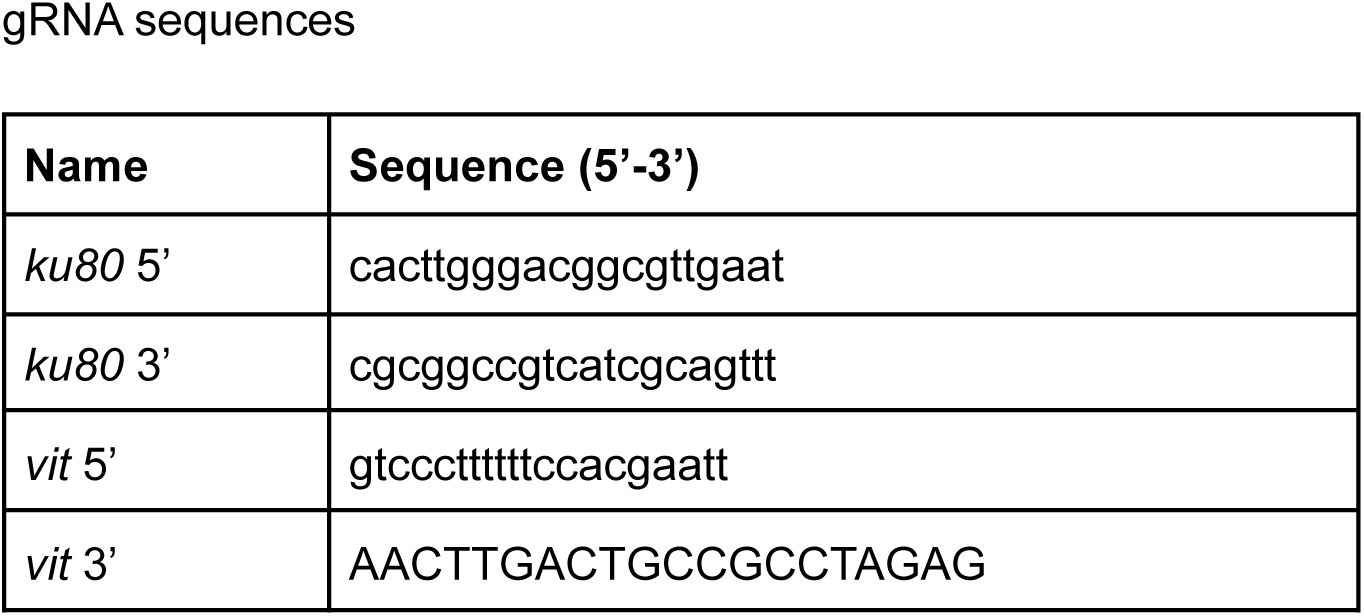
A list of the gRNA sequences used in this publication

### ΔVIT parasites are hypersensitive to excess iron and have lower cellular iron levels

In yeast and *Plasmodium*, deletion of CCC1 or *Pb*VIT leads to hypersensitivity to excess iron (Li et al., 2001; Slavic et al., 2016). To examine this in *T. gondii*, we performed a plaque assay under increasing concentrations of ferric ammonium chloride (FAC). We found that ΔVIT parasites were noticeably more sensitive to excess FAC, forming fewer and much smaller plaques than the parental line (**Fig. 2A**). To confirm this result, we performed a competition assay under standard conditions and with excess FAC. To serve as controls, two new lines were constructed in RhΔHX, where the coding region of Ku80 was replaced with a cassette expressing mNeonGreen (ΔKu80::mNeon, called mNeon) or tdTomato (ΔKu80::tdTomato) (**Fig. S1A**), resulting in two lines which expressed fluorescent proteins in place of the endogenous *ku80* gene. We confirmed that these lines were fluorescent using flow cytometry and the loss of the *ku80* gene by PCR (**Fig. S1B**). To assess the fitness defect of the ΔVIT strain, ΔVIT or mNeon parasites were mixed with tdTomato in an equal ratio and untreated or treated with 200 μM FAC. The proportion of green parasites (either mNeon or ΔVIT) in the population was assessed at each passage by flow cytometry. The proportion of mNeon parasites did not change over the experiment, however, the ΔVIT line was significantly *(p* = 0.002, *t* test, Holm-Sidak corrected) outcompeted by three days post infection (**Fig. 2B**). Addition of FAC exacerbated this phenotype, ΔVIT parasites were significantly *(p* = 0.001, *t* test, Holm-Sidak corrected) outcompeted by two days post infection in the presence of excess iron and were almost undetectable by four days post infection. This shows that VIT is required for growth under normal conditions, and lack of VIT sensitises parasites to exogenous iron. To confirm this, we constructed a second ΔVIT strain by replacing the *vit* gene with a DHFR cassette (**Fig. S2A**), named ΔVIT::DHFR_TS_. We confirmed integration by PCR (**Fig. S2B**) and confirmed these parasites were also hypersensitive to excess iron by plaque assay (**Fig. S2C and D**).

**Figure 2.**
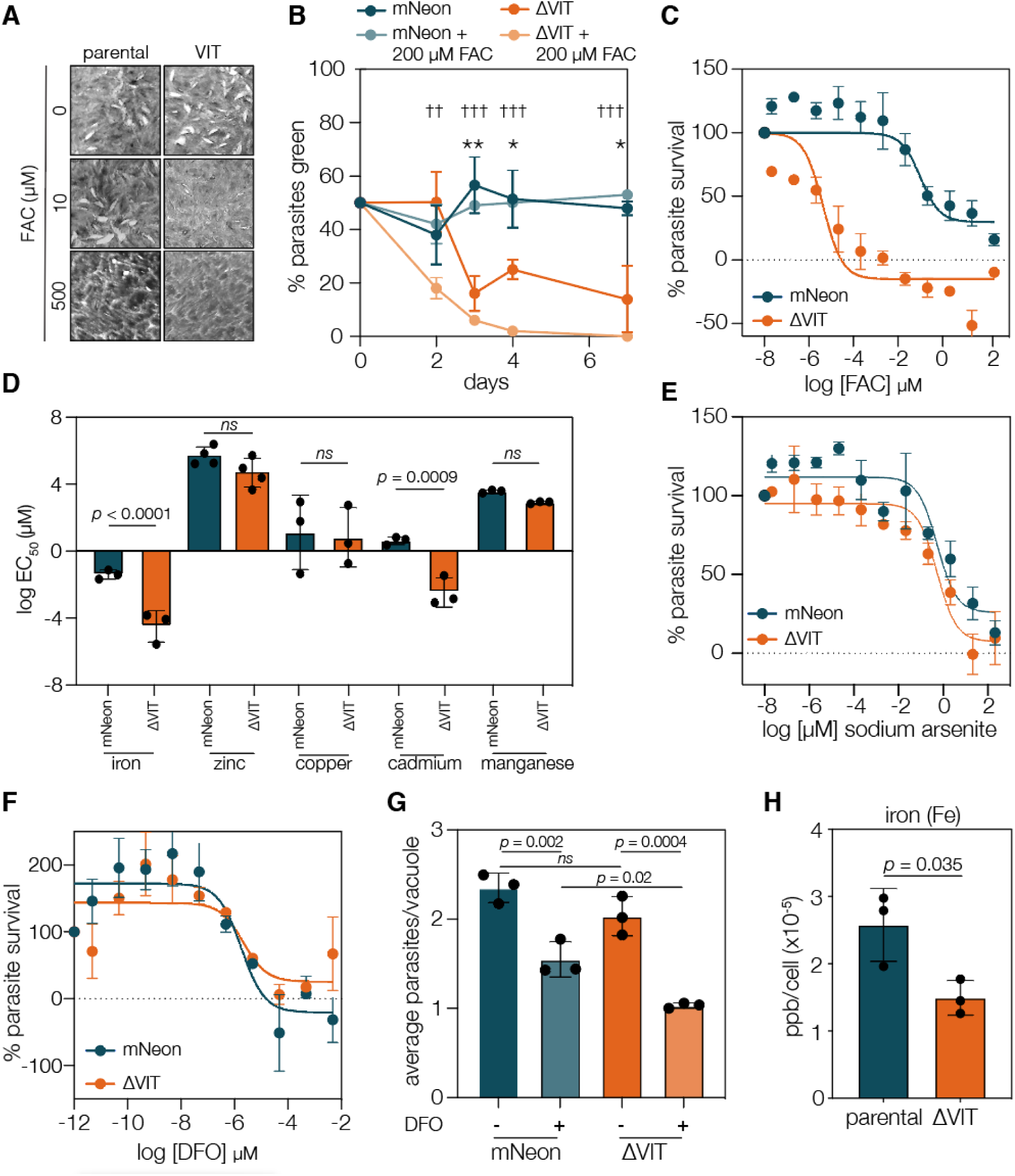
ΔVIT parasites are more susceptible to iron overload. **A**. Plaque assay performed under indicated concentrations of iron (ferric ammonium chloride, FAC). ΔVIT parasites showed increased susceptibility to exogenous iron and made smaller plaques. **B**. ΔKu80::tdTomato were mixed with ΔKu80::mNeon and ΔVIT parasites in a 1:1 ratio and untreated or treated with 200 μM FAC. ΔVIT parasites were significantly outcompeted by three days post infection. Points are the mean of four independent experiments ± SD. * *p* < 0.01, ** *p* < 0.005, t test between ΔKu80::mNeon and ΔVIT, †† *p* < 0.005, ††† *p* < 0.0001, t test between ΔKu80::mNeon + 200 μM FAC and ΔVIT + 200 μM FAC, corrected with Holm-Sidak **C**. ΔVIT parasites were significantly more sensitive to excess FAC than the mNeon line. mNeon Points are the mean of three independent experiments, ± SEM. **D**. Graph showing mean EC_50_ for iron, zinc, copper, cadmium, and manganese for mNeon and ΔVIT parasites, each point represents an individual experiment performed in triplicate. Bars at the mean of experiments, ± SD. *p* values from extra sum of square F test. **E**. Treatment of mNeon and ΔVIT parasites with sodium arsenite (Ars), a ROS generator, showed no difference in sensitivity between the two lines. Points are the mean of at least three independent experiments, ± SEM. **F**. ΔVIT parasites did not show any significant change in the EC_50_ upon treatment with the iron chelator DFO. Results are the mean of three experiments, ± SEM. **F**. mNeon and ΔVIT parasites were allowed to invade HFF cells untreated, or pretreated with DFO for 24 h. At 14 h post invasion, average parasite/vacuole were quantified. Results mean ± SD of three independent experiments, at least 100 vacuoles counted/experiment. **G**. ICP-MS of parental and ΔVIT::DHFR_TS_ parasites. Bars are at the mean ± SD of three independent experiments, *p* value from *t* test.

To quantify this hypersensitivity, we infected host cells in 96 well plates with mNeon or ΔVIT parasites and treated them with increasing concentrations of FAC for four days before quantifying parasite fluorescence (**Fig. 2C**). The ΔVIT parasites were significantly (*p* < 0.0001, extra sum of squares F-test) more sensitive to FAC treatment, with an EC_50_ of 0.004 μM (95% C. I. 0.001 to 0.018 μM), compared to the mNeon EC_50_ of 102 μM (95% C.I. 20 to 490 μM), an increase in sensitivity of around 25,000-fold.

VIT homologs from various species have been shown to transport other metals, including manganese (Li et al., 2001) and zinc (Zhang et al., 2012). To test if deletion of VIT altered the sensitivity of *T. gondii t*o metals other than iron, we treated parasites as above with increasing concentrations of zinc, copper, cadmium, and manganese and determined the EC_50_. We saw that the absence of VIT did not affect the response to treatment with excess zinc, copper, and manganese (extra sum of squares F-test, *p* values > 0.05) (**Fig. 2D and S3A-D**). While this does not prove that VIT is unable to transport these substrates, it does show that VIT is not required for the detoxification of the selected metals.

However, we did see a significant shift in the sensitivity to the heavy metal cadmium (EC_50_ of parental line 3.41 μM (95% C.I. 1.63 to 6.34 μM), EC_50_ of ΔVIT 0.04 μM (95% C.I. 0.008 to 0.149 μM), *p* = 0.0009, extra sum of squares F-test) (**Fig. 2D and S3C**). Cadmium has complex effects on mammalian cells and tissues (Templeton and Liu, 2010) and there is evidence that cadmium can lead to increased ROS production (Shaikh et al., 1999). To determine if the ΔVIT parasites were more susceptible to general increases in oxidative stress, we treated the parasites with sodium arsenite (Ars) which induces oxidative stress through a number of mechanisms (Hu et al., 2020) and has been successfully used in *T. gondii* to induce oxidative stress (Augusto et al., 2021). The concentrations of Ars used did not affect host cell viability as measured by Alamar blue assay (**Fig. S3E**). We saw no significant change in the sensitivity between parental line and the ΔVIT parasites to Ars treatment (**Fig. 2E**), demonstrating that the effect of VIT on cadmium toxicity is somewhat specific, and that ΔVIT parasites do not have a general sensitivity to oxidative stress. VIT from the distantly related diatom *Phaeodactylum tricornutum* is transcriptionally regulated by cadmium (Brembu et al., 2011) where the authors suggest that Cd^2+^ may also be stored in a vacuole. This has not previously been examined in the Apicomplexa, however our results here may suggest some link between iron and cadmium detoxification in *T. gondii*.

In *Plasmodium*, deletion of VIT also led to a change in the sensitivity to the iron chelator deferoxamine (DFO) (Slavic et al., 2016). However, in *T. gondii* we saw no change in the ability of the parasites to replicate in the presence of increasing concentrations of iron chelator (**Fig. 2E**) (EC_50_ of mNeon 160 μM (95% C.I. 65.9 to 362.2 μM) and ΔVIT 101.3 μM (95 % C.I. 44.9 to 233.5 μM), *p* > 0.05, extra sum of squares F-test) over the four days of the experiment.

If VIT is responsible for storing iron, it is possible that in its absence, parasites would be disadvantaged in replication when moving to an iron-poor environment. To simulate this, we pretreated host cells with DFO for 24 h to deplete iron, before adding mNeon or ΔVIT parasites and fixed after only 14 h post invasion. We then quantified the average number of parasites/vacuole. We found that pretreatment with DFO significantly reduced the average parasites/vacuole in both the mNeon and the ΔVIT parasite lines (**Fig. 2F**), demonstrating the importance of iron to parasite replication. Although we do see a replication defect in the ΔVIT strain (**Fig. 1C**), at this early point in infection it was not significant. However, in an iron-poor environment there was a significant decrease in the average parasite/vacuole in the ΔVIT strain compared to the mNeon strain, suggesting that the potential disruption of iron storage inhibits replication in an iron-depleted environment.

The above results suggested a loss of stored iron. To assess if iron levels changed in the ΔVIT::DHFR_TS_ parasites, we quantified total parasite-associated iron using inductively coupled plasma-mass spectrometry (ICP-MS). Using this method, we found a mean of 2.6 × 10^−5^ ppb of Fe/parasite in the parental parasite line (**Fig. 2G**). In the absence of VIT, we saw a significant (*p* = 0.035, *t* test) reduction in iron to approximately 1.5 × 10^−5^ ppb/parasite, approximately 60% of the parental level. As a control, we saw no change in the levels of zinc between the parasite lines (**Fig. S2E**). A reduction in total iron has been observed before upon disruption of vacuolar iron transporters in yeast and plants (Kim et al., 2006; Li et al., 2001), and confirms the role of VIT in *T. gondii* iron storage.

### VIT has a dynamic localisation throughout the lytic cycle

In *Plasmodium*, VIT colocalized with markers of the endoplasmic reticulum (ER) throughout the parasite’s lifecycle (Slavic et al., 2016). To determine the localisation of VIT in *T. gondii*, we first overexpressed VIT with a C-terminal Ty tag from the TUB promotor. Transient overexpression of VIT-Ty resulted in localisation of several foci within the cell with some expression at the parasite periphery (**Fig. 3A**). The central foci did not colocalise with the apicoplast marker HSP60 but did show a degree of overlap with the marker of the vacuolar associated compartment (VAC), cathepsin protease L (CPL). However, overexpression of VIT may be toxic as we did not see any parasites past the two-to-four cell stage. To determine the native localisation, we endogenously tagged VIT using 3xHA tags at the C-terminus (**Fig. 3B**). We also saw that endogenously tagged VIT-HA colocalized with the overexpressed VIT-Ty (**Fig. S4**) although we saw less expression at the periphery of the parasite. We also confirmed the expression of a single band of VIT-HA at the expected size (35 kDa) by western blot (**Fig. 3C**). Upon IFA, we observed VIT-HA expression as a single point in extracellular parasites. This localisation was maintained at 1 h post infection, but between 1 and 6 h post infection this point appeared to fragment and by 24 h post infection VIT-HA signal was seen in multiple small structures throughout the cytoplasm of the parasite (**Fig. 3D**). We quantified this fragmentation using an automated ImageJ macro and R code and saw a significant increase (*p* < 0.0001, one way ANOVA with Tukey correction) in the number of VIT-HA foci per parasite between 1 and 6 h post infection (**Fig. 3E**). We also assessed the amount of VIT-HA by western blot and found that the change in localisation was not associated with a significant change in protein level (**Fig. 3F and G**). Dynamic localisation through the lytic cycle has previously been observed in proteins localizing to the vacuolar associated compartment (VAC) (Thornton et al., 2019; Warring et al., 2014) and as we saw some overlap with CPL with the overexpressed VIT-Ty, we tested if VIT-HA colocalised with CPL. However, we found that VIT-HA did not colocalize with an antibody against CPL at 1 h post infection (**Fig. 3H**). Interestingly, we did observe frequent colocalization between VIT and CPL when VIT was tagged with a Myc epitope (**Fig. 3H**). If VIT localises, even transiently, to the VAC it is possible that the VAC is the site of iron storage in the cell. To determine how the VAC responded to excess iron, we treated parasites with high levels of iron (2.5 mM FAC) to be sure of overwhelming any host cell buffering effects and determined the area of the VAC in an unbiased manner using an ImageJ macro and the luminal VAC marker CPL. We found that the total area of CPL signal significantly increased (**Fig. 3I**) when parental parasites were treated with high exogenous iron (*p* = 0.0008, one way ANOVA with Dunnett’s correction). We saw no change in VAC area between the parental and ΔVIT parasites, but interestingly when we treated ΔVIT parasites with excess iron we did not see a change in the VAC area. These results suggest that VAC morphology is altered by excess iron through the actions of VIT, providing supporting evidence that VIT is localized to this compartment in *T. gondii*.

**Figure 3.**
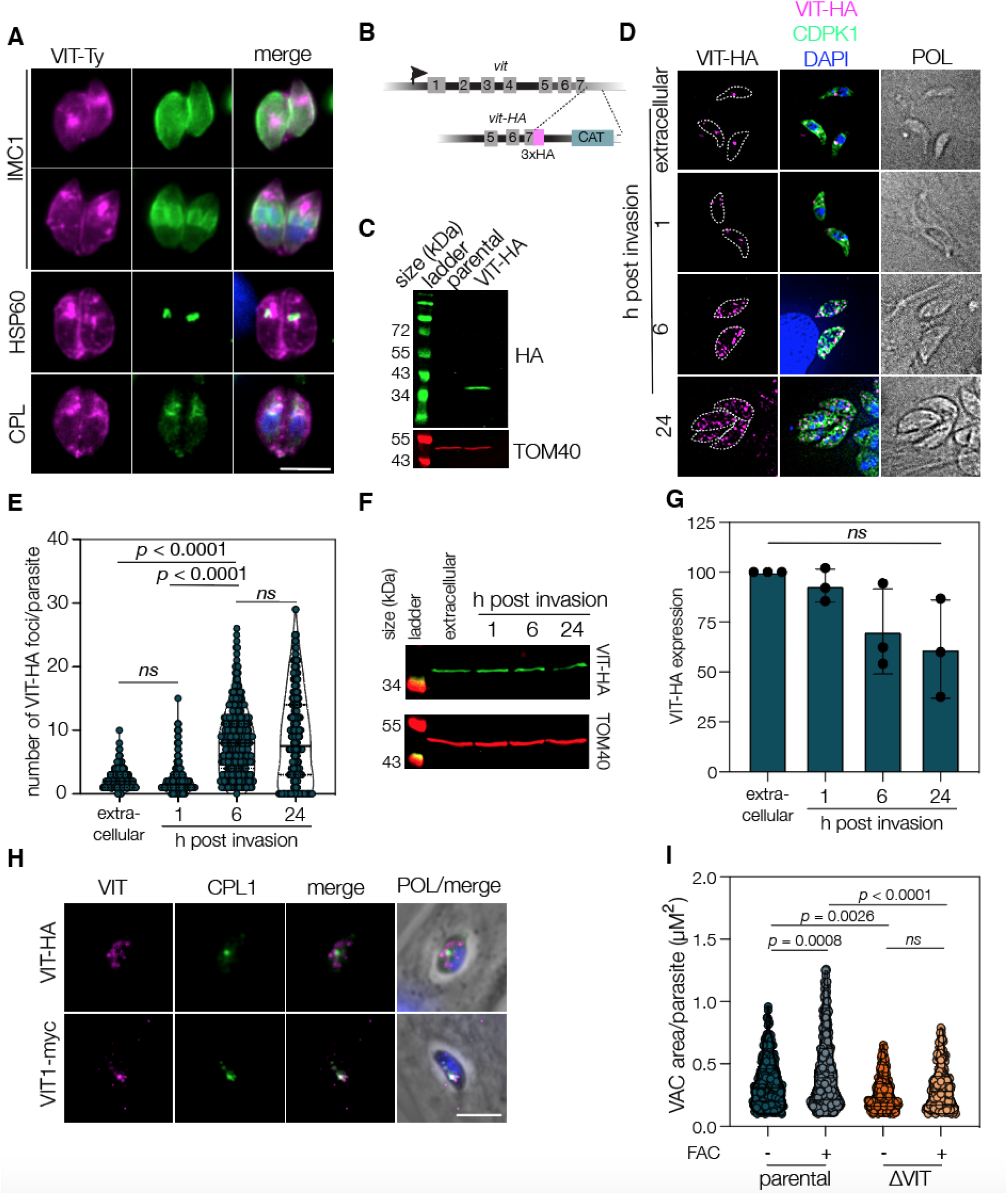
VIT has a dynamic localisation which alters through the lytic cycle. **A**. Transient overexpression of VIT-Ty in the parental parasite line. VIT-Ty (magenta) showed colocalisation with the periphery of the parasite (IMC1) and the VAC marker CPL, but not the apicoplast marker HSP60 (all green). Scale bar 5 μm. **B**. VIT was endogenously tagged with 3xHA tags at the C-terminus. **C**. Western blot against HA epitope demonstrated a single band at around 35 kDa. **D**. IFA of VIT-HA showing a dynamic localisation through the lytic cycle, VIT-HA exists at a single point in extracellular and 1 h post invasion parasites before fragmenting at 6 - 24 h post invasion to a number of small foci throughout the parasite. Scale bar 5 μm. **E**. Violin plot of number of foci/parasite from three independent experiments, 100 parasites quantified/experiment. Bars at mean and quartiles. *p* values from one way ANOVA with Holm-Sidak correction. **F**. Western blot showing VIT-HA at indicated time points, TOM40 used as a loading control. **G**. Quantification of VIT-HA levels from three independent experiments, ns - no significant change. **H**. VIT-HA does not co-localise with the VAC marker CPL at 1 h post invasion, however endogenously tagged VIT-myc did show a degree of colocalisation. Scale bar 5 μm. **I**. mNeon and ΔVIT parasites were treated for 1 h with excess FAC (2.5 mM) and the area of the VAC (as assessed by CPL staining) was quantified. Results of two independent experiments, bar at mean ± SD

### VIT expression is regulated by iron availability

The expression, localisation and activity of transporters are frequently regulated by the availability of the substrate. This has been well established in model organisms (Conte and Walker, 2011; Gao and Dubos, 2021; Lee et al., 2020; Sorribes-Dauden et al., 2020) and was recently reported for the arginine transporter ApiAT1 in *T. gondii* (Rajendran et al., 2021). To determine if iron affects expression of VIT we performed qRT-PCR on the *vit* gene. Unexpectedly, we found that transcript levels were significantly decreased (*p* = 0.007, one sample *t* test) (**Fig. 4A**) at 24 h upon treatment with excess iron. This is the opposite to the situation seen in yeast, where excess iron upregulates CCC1 (Li et al., 2008). In contrast, removal of iron by DFO treatment did not lead to a change in VIT RNA. As transcript and protein levels do not always correlate, we examined changes at the protein level by western blotting using VIT-HA. We found that treatment with FAC also led to a variable but significant (*p* = 0.043, one sample *t* test) decrease in VIT protein levels (**Fig. 4B and C)**, however interestingly, removal of iron by DFO also led to a decrease (*p* = 0.032, one sample *t* test) in VIT-HA levels despite no change in RNA levels. As well as regulation at the mRNA and protein level, the function of transporters can also be regulated by changes in localisation (Lee et al., 2020). We assessed the localisation of VIT-HA upon changes in iron levels. As described above, we quantified the number of VIT-foci after treatment with FAC or DFO at 6 and 24 h post infection. At 6 h post infection we saw a significant (*p* = 0.0004, one way ANOVA with Tukey) decrease in the number of foci/parasite upon treatment with DFO but not FAC (**Fig. 4D**). By 24 h post infection we saw a significant decrease in foci number under both conditions (p < 0.0001, one way ANOVA with Tukey) (**Fig. 4E**). This may show a different response to high or low iron, or may reflect the speed at which the treatments take effect within the cells. In either case, changes in iron levels appear to alter the transcription, protein level and localisation of VIT within the parasite.

**Figure 4.**
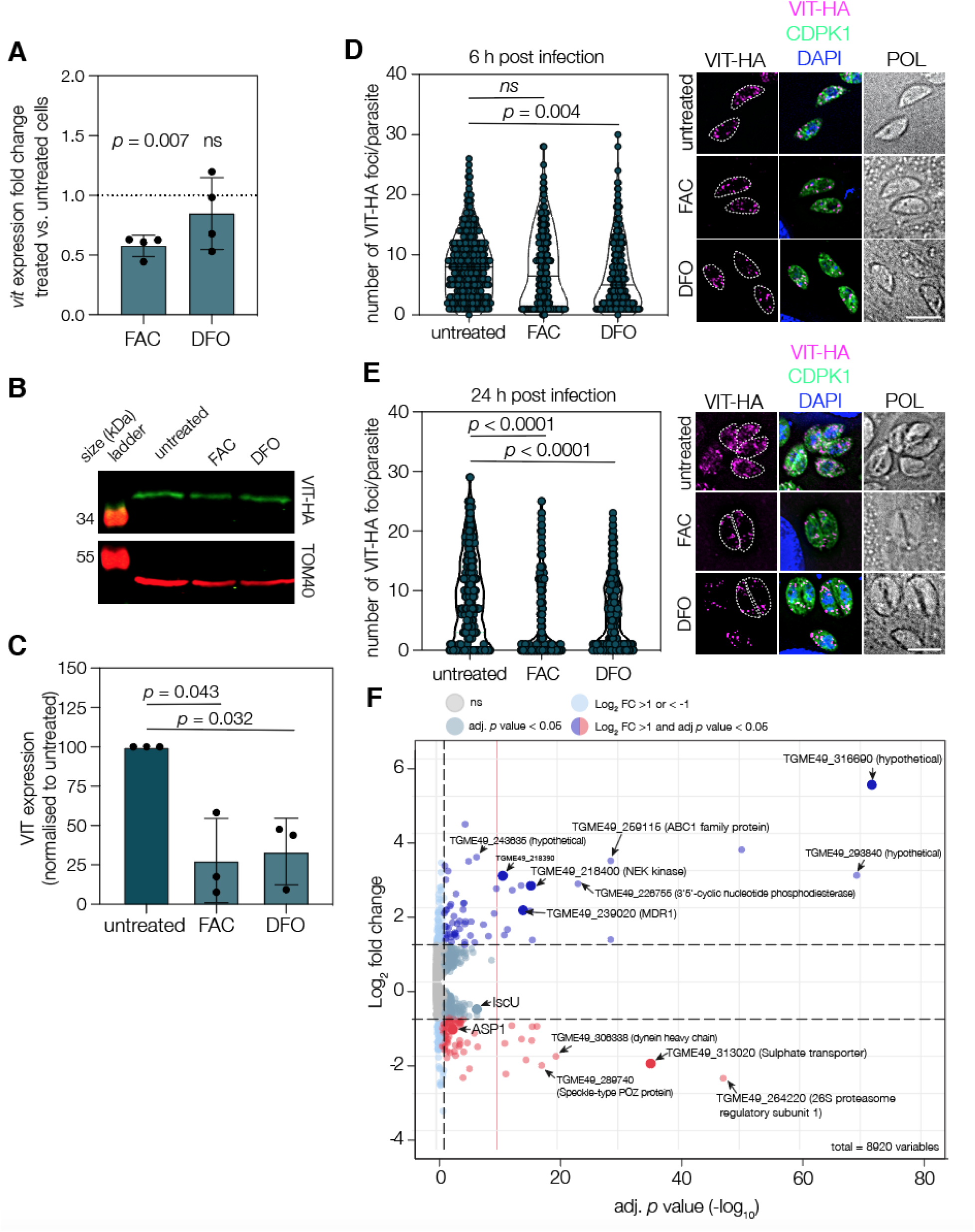
VIT expression is regulated by the changes in iron levels. **A**. qRT PCR on *vit* transcripts after treatment with FAC or DFO, normalised to actin. Points represent 4 independent experiments, bars at mean ± SD. *p* values from one sample *t*-test. **B**. Western blot showing levels of VIT-HA after 24 h treatment with 100 μM DFO or 5 mM FAC. TOM40 used as a loading control. **C**. Quantification of VIT-HA levels from three independent experiments, *p* values from one sample *t* test. **D**. VIT-HA foci at 6 h post invasion after treatment with FAC or DFO. There was no significant change upon FAC treatment, however there was a significant decrease in the number of foci upon DFO treatment. Results from three independent experiments, at least 100 parasites/replicate. *p* values from one way ANOVA with Tukey correction. **E**. As above, but quantified at 24 h post invasion. There was a significant decrease in the number of foci after treatment with both FAC and DFO, *p* values from one way ANOVA with Tukey correction. Results from three independent experiments, at least 100 parasites/replicate. **F**. Volcano plot from RNAseq data comparing the parental line to ΔVIT. See text for more details.

### Transcriptional response to the absence of VIT

The finding that VIT is transcriptionally regulated by iron led us to examine how the transcriptome changes in the absence of VIT and correct iron storage. RNAseq was performed on the parental and ΔVIT strains in triplicate. Under these conditions, 70 genes were downregulated upon deletion of VIT (log_2_ fold change (LFC) < -1, adjusted p value (Padj) < 0.05) and 62 were upregulated (log_2_ fold change > 1, adjusted p value < 0.05) under standard growth conditions (**Fig. 4F, Table S1**). One gene that was upregulated upon deletion of VIT was an ABC1 family protein (TGME49_259115), a similar protein in plants is upregulated by Cd^2+^ toxicity and is believed to act in oxidative stress signalling (Jasinski et al., 2008). We also saw significant upregulation of TGME49_313020, a predicted plasma membrane sulphate transporter raising the possibility that iron and sulfur transport are linked in *T. gondii*. We also saw significant upregulation of the potential plasma membrane ABC-type transporter (TGME49_239020). This gene had strong structural homology (E = 7.1 e^-135^, HHPRED (Söding et al., 2005)) to multidrug efflux transporters and is named multidrug resistance protein 1 (MDR1) in *Plasmodium* (Koenderink et al., 2010). Interestingly, work in mammalian cancer cells has shown that removal of iron led to a decrease in MDR1 levels (Fang et al., 2010) and disrupting iron storage through increased expression of ferritin led to an increase in MDR expression (Epsztejn et al., 1999). Upregulation of this transporter may thus be an adaptation by the cell to attempt to remove excess iron from the parasite, in the absence of correct storage.

There are two well studied pathways involving iron in *T. gondii*, the heme biosynthesis pathway and iron-sulfur cluster biogenesis. The contribution of VIT-based iron storage to these pathways is currently unknown. We examined the transcriptional changes in the heme biosynthesis pathways (**Fig. S5A**) and found there was a general upregulation of several components of the pathway upon VIT deletion, most significantly PBGD. The only gene in this pathway which directly interacts with iron — ferrochelatase — was not altered, however there was a significant upregulation in a cytochrome c heme lyase, one of the most important destinations for heme in the parasite. We also saw downregulation of the oxygen-independent coproporphyrinogen-III oxidase, however this gene is dispensable to the parasite and its role (if any) is not well understood (Krishnan et al., 2020). *Toxoplasma* encodes three iron sulfur cluster biogenesis pathways, localised to the cytosol, mitochondrion and apicoplast. Interestingly, we saw opposing changes in the mitochondrial ISC pathway, with downregulation of the scaffold IscU and upregulation of the acceptor IscA (**Fig. S5B**). In both mammalian and yeast cells, IscU expression is iron dependent, suggesting that the altered iron storage in the ΔVIT parasites is altering iron-dependent processes within the cell (Garland et al., 1999; Tong and Rouault, 2006).

Ribonucleoside-diphosphate reductases (RNR) are enzymes involved in DNA repair, which contain ferrous iron. We saw significant downregulation of RNR large chain in the absence of VIT. This has been observed in yeast where RNR subunits are under the control of the iron-dependent mRNA decay factor Cth1/2 and are downregulated upon iron deficiency as the cell prioritises certain pathways for iron (Sanvisens et al., 2011). Other essential genes containing iron, including catalase, SOD2 and SOD3 and prolyl hydroxylases were not transcriptionally affected in the absence of VIT. This provides the first hints that in the absence of correct iron storage, *T. gondii* also has an iron prioritisation programme, however future work will be needed to understand this intriguing possibility. Overall, we saw significant changes in transcription upon deletion of VIT, with changes in transcription of genes and pathways known to require iron.

### Iron overload leads to ROS accumulation in the absence of VIT

Ferrous iron can lead to the production of reactive oxygen species (ROS) though the Fenton reaction (Imlay et al., 1988) and loss of CCC1 in yeast is associated with increased oxidative stress (de Llanos et al., 2016). To determine if absence of VIT affected ROS accumulation in *T. gondii*, we stained parasites with CellROX Deep Red, which fluoresces in the presence of ROS, and quantified fluorescence by flow cytometry. In the parental line, treatment with very high concentrations of FAC (2 mM) did not significantly increase the ROS levels, suggesting the presence of effective mechanisms for controlling iron toxicity in these cells. Although there was no increase in ROS levels under normal conditions in the ΔVIT strain, upon FAC treatment ROS levels were significantly (*p* = 0.004, one way ANOVA with Sidak’s correction) higher than in the parental line (**Fig. 5A**). CellROX is localised in the cytosol of the parasites, to determine if levels in the mitochondrion were changed, we made use of the mitochondrially-localised MitoNeoD (Shchepinova et al., 2017) which increases in fluorescence in the presence of mitochondrial superoxide. However, we did not see any significant changes in the levels of MitoNeoD fluorescence in the ΔVIT parasites (**Fig. S6A)** in the presence or absence of exogenous iron, suggesting that the cytosol is the major location for ROS production upon excess iron treatment.

**Figure 5.**
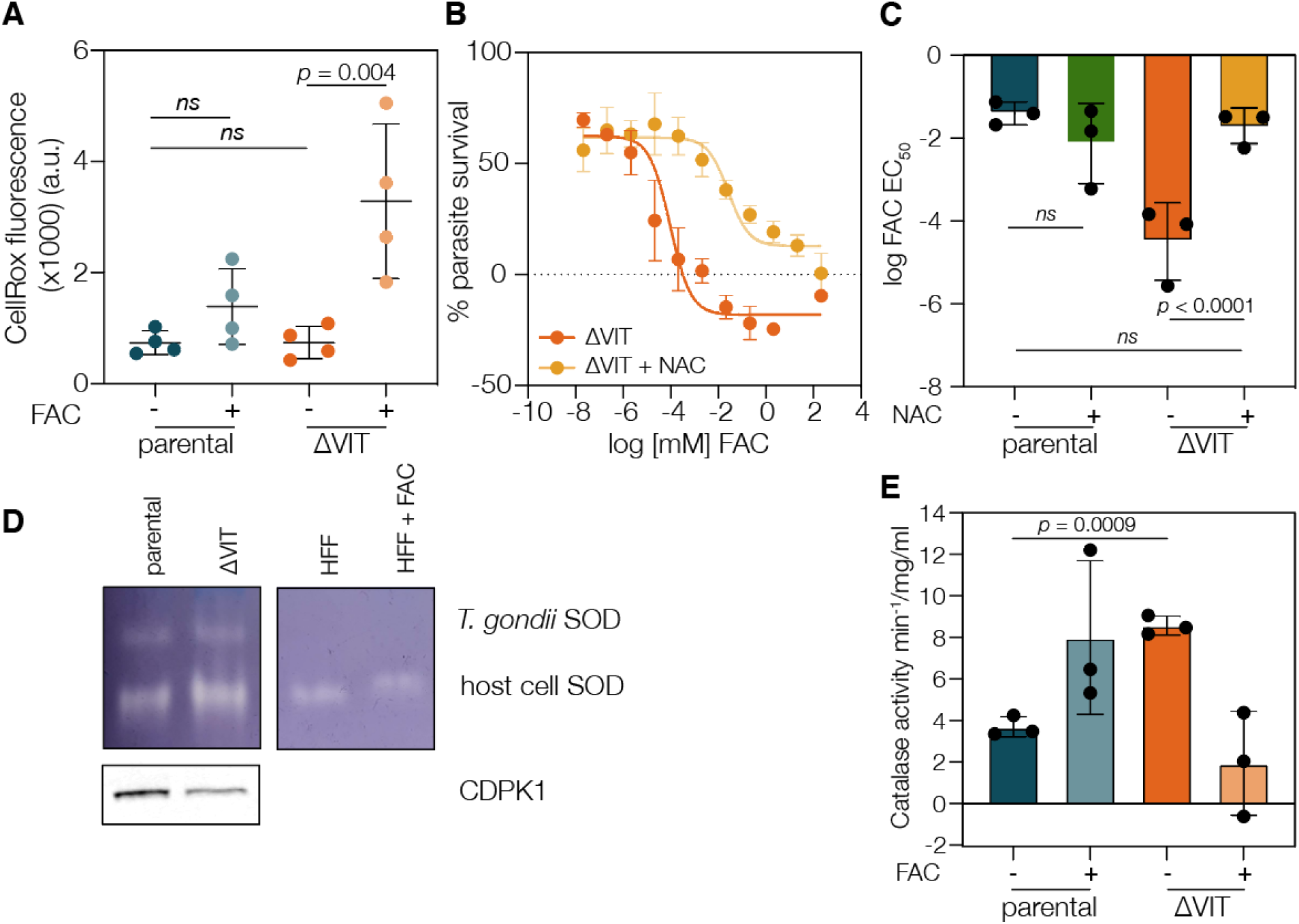
Iron overload in absence of VIT leads to ROS accumulation. **A**. Parental and ΔVIT parasites were treated with FAC overnight, and CellROX Deep Red fluorescence quantified using flow cytometry. Iron treatment of the parental line had no significant effect on CellROX fluorescence, however in the ΔVIT line CellROX was significantly (*p* = 0.004, one way ANOVA with Sidak correction) increased upon iron overload. Each point represents geometric mean fluorescence of over 10,000 cells from a single biological replicate, ns - not significant. Bars are at mean, ± SD. **B**. Treatment with the NAC (5 mM) rescued ΔVIT parasite sensitivity to FAC. Points are the mean of at least three independent experiments, ± SEM. **C**. Graph showing the mean EC_50_ for FAC of mNeon and ΔVIT parasites, with or without NAC treatment. Each point represents an individual experiment performed in triplicate, bars at the mean of three experiments, ± SD. *p* values from extra sum of square F test. **D**. Superoxide dismutase activity was examined by clear bands on a NBT stained native gel. Parasite SOD was clearly distinguishable from host cell enzymes as migrated at a different rate. A western blot against CDPK was used as a parasite loading control. Results representative of at least two independent experiments. **E**. ΔVIT parasites had significantly higher catalase activity compared to the parental line. Each point represents an independent experiment performed in triplicate, bar at mean ± SD. *p* value from one way ANOVA with Sidak correction.

To determine if the increase in ROS induced by FAC in the ΔVIT line is responsible for the increased sensitivity, we treated the parasites with the ROS scavenger N-acetyl-cysteine (NAC) (Charvat and Arrizabalaga, 2016). Upon 5 mM NAC treatment, the FAC EC_50_ for the parental line increased slightly (from 88.28 μM, 95 % C.I. 24.4 to 307 μM to 109.7 μM, 95 % C.I. 15 to 136 μM), but not significantly (**Fig. 5B and C**). However, scavenging of ROS by NAC led to a significant (*p* < 0.0001, extra sum of square F test) protective effect (EC_50_ 3.8 μM, 95 % C.I. 0.08 to 5.5. μM, compared to 0.018 μM, 95 % C. I. 0.0021 to 0.1 μM) to excess iron in the ΔVIT parasites (**Fig. 5B and C**). This demonstrates that a significant driver of the increased sensitivity of the ΔVIT parasites to FAC is the increased production of ROS.

Although the ΔVIT parasites have a small growth defect *in vitro*, we were intrigued to see that there was no increase in ROS under normal conditions. The parasite has various lines of defense against ROS, including thioredoxins, several SODs and a cytosolically localised catalase (Alves et al., 2021; Kwok et al., 2004; Xue et al., 2017). Although the RNAseq did not show any changes in SOD, SOD2 (**Fig. S6B**) or catalase transcription (**Fig. S6C**), we decided to investigate the activity of these enzymes. We investigated SOD activity using a native page in gel activity assay, however saw no large difference between the parental and ΔVIT parasites (**Fig. 5D**), CDPK1 was used as a parasite loading control. *T. gondii* encodes for three SOD enzymes, two of which are expressed and localised to the cytosol and mitochondria (Kwok et al., 2004; Sibley et al., 1986) and we were only able to distinguish one from the host cell SOD enzymes. Given this, we cannot exclude the possibility that SOD activity overall did change, but we were unable to visualise this. We also assessed the enzyme activity of catalase using a colorimetric assay (Hadwan, 2018; Portes et al., 2015). We found that deletion of VIT led to a significantly (*p* = 0.0009, Welch’s one-way ANOVA with Dunnett’s correction) increased catalase activity compared to the parental line under normal growth conditions (**Fig. 5E**). This suggests that to maintain growth, ΔVIT parasites require higher catalase activity. This is despite the lower amounts of parasite-associated iron, potentially due to the mislocalisation of iron in these parasites.

### VIT is required for virulence *in vivo*

The increased activity of catalase and the reduced growth in cells suggested that VIT has an important role in survival of *T. gondii in vitro*. To determine if VIT had a role in causing disease, we infected five Swiss Webster mice peritoneally with either 20 or 100 tachyzoites of the parental or ΔVIT::DHFR_TS_ parasites and assessed mouse survival over time (**Fig. 6A**). Deletion of VIT led to significantly increased survival after infection with both 20 (*p* < 0.0001, log-rank test) and 100 parasites (*p* = 0.0027, log-rank test) demonstrating that VIT has an important role in virulence *in vivo*.

**Figure 6.**
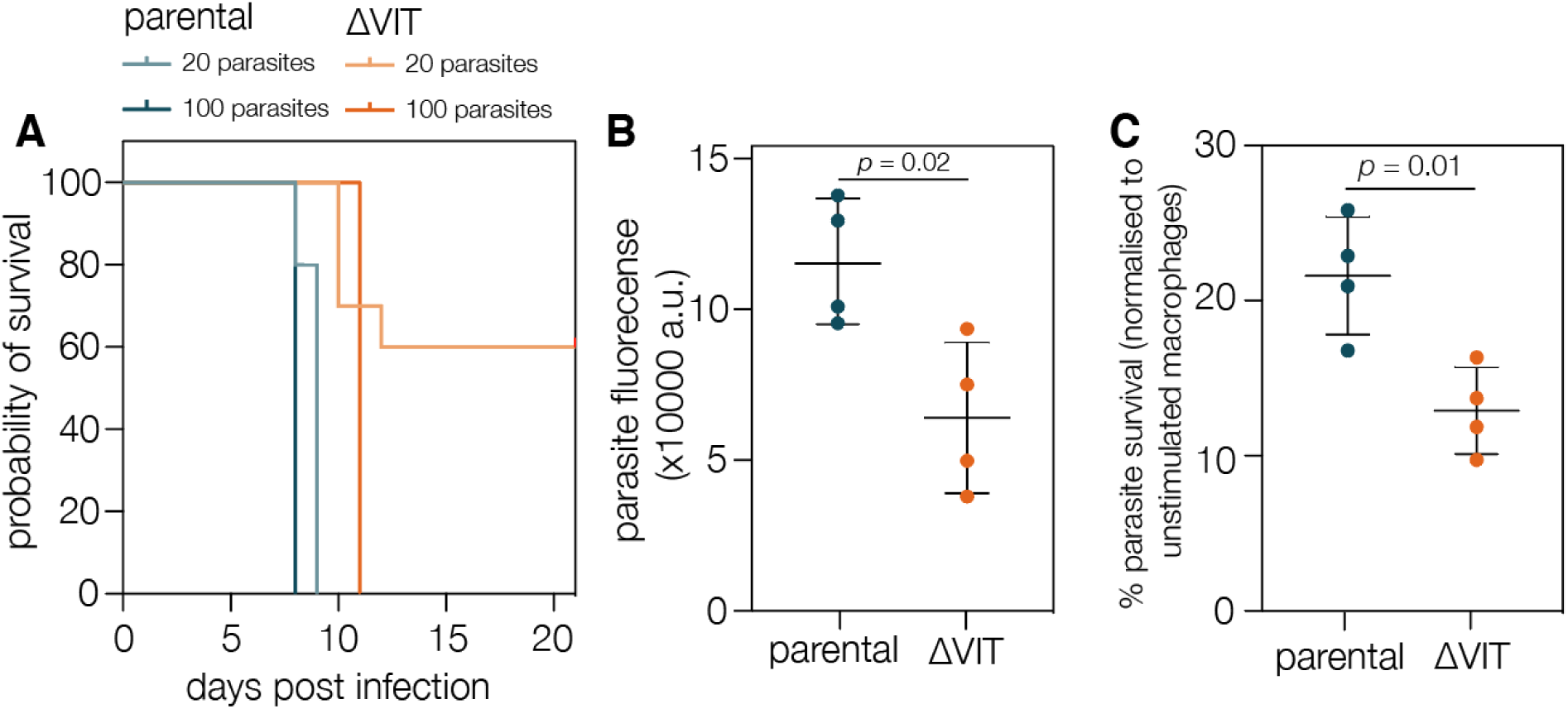
VIT is required for parasite survival in macrophages and for pathogenesis *in vivo*. **A.** 10 mice were infected with 20 or 100 tachyzoites of the indicated strain and survival monitored over the course of the experiment. **B**. mNeon and ΔVIT parasites were used to infect macrophages at an MOI of 5 and fluorescence (arbitrary units) monitored after 72 h. Results are mean of four independent experiments, performed in triplicate, ± SD. **C**. After normalisation of the fluorescence in the unstimulated macrophages (from B) there was a significant decrease in survival of the ΔVIT parasites in stimulated (IFNy/LPS) macrophages. Results are the mean of four independent experiments, performed in triplicate, ± SD.

Given the relatively small growth defect seen under normal growth conditions, we were surprised by the magnitude of the effect *in vivo*. In order to cause disease, *T. gondii* must evade the host immune system and colonise various tissues. One important bottleneck is the ability of the parasite to survive in activated phagocytic cells (Lykens et al., 2010; Wang et al., 2020). To determine if loss of VIT affected the parasite’s ability to survive, we prestimulated the murine monocytic cell line RAW 264.3 with LPS and interferon-*γ* (IFN*γ*) before infecting them at an MOI of 5 for 48 h. We quantified fluorescence and normalized levels in stimulated macrophages to unstimulated cells. ΔVIT parasites demonstrated less fluorescence even in unstimulated macrophages (**Fig. 6B**), but after normalisation we saw that ΔVIT parasites had a significantly (*p* = 0.03, *t* test) reduced ability to survive in activated macrophages, with a survival of around 12% compared to 21% survival in the mNeon line (**Fig. 6C**). Although virulence is a complex process, it is likely that the reduced survival of the ΔVIT parasites in macrophages contributed to their inability to cause severe disease in mice

## Discussion

Iron plays a central role in metabolism; however, its potential toxicity means that transport and storage are tightly regulated. Here, we have investigated VIT, the first characterised iron transporter in *T. gondii*, and shown that it has a role in the detoxification of excess iron and in survival of the parasite. Deletion of VIT leads to parasite hypersensitivity to iron and a decrease in stored iron within the cells. We find that VIT localises to a vesicular compartment within the cell, which changes in distribution throughout the lytic cycle. VIT is regulated at the transcript and protein level by iron availability and the absence of VIT-mediated iron detoxification leads to transcriptional changes in the cell, including in pathways and genes known to rely on iron. We show that in the absence of VIT parasite hypersensitivity to excess iron can be reversed by scavenging of excess ROS, suggesting that iron-generated ROS is the major cause of parasite death. Finally, we show that VIT is required for survival in activated macrophages, and in pathogenesis *in vivo*, confirming that iron detoxification plays an important role in the parasite.

Lack of VIT causes hypersensitivity (over 20,000 fold increased sensitivity) to excess iron in *T. gondii*. This dramatic phenotype is mostly specific to iron, as no change in the sensitivity to other metals was seen (with the exception of cadmium). This supports the previous data from *Plasmodium* that the Apicomplexan VIT is a selective iron transporter (Slavic et al., 2016), unlike the homologues from rice or yeast which have been shown to transport zinc and manganese in addition to iron (Li et al., 2001; Zhang et al., 2012). In further support, we saw no changes in parasite-associated zinc upon VIT deletion. In *T. gondii*, lack of VIT was also associated with decreased parasite-associated iron and a decreased ability to replicate in acute iron deprivation. These results are congruent with studies on the VIT transporters of yeast and plants which also showed that deletion of CCC1/VIT was associated with reduced cellular iron (Gollhofer et al., 2014; Li et al., 2001). However, in *Plasmodium*, deletion of VIT was associated with an increase in chelatable, or labile, iron in the cells (Slavic et al., 2016). We were not able to assess the labile iron in our cells using a similar method. However, our results may well be consistent with data from *Plasmodium*, it is possible that even though the total parasite-associated iron is reduced, the labile iron in the cytoplasm increases. This would also make sense in the context of the increased cytosolic ROS upon iron treatment. Alternatively, it is possible that VIT, and iron storage in general, have different roles within the parasite species, which may require further studies to appreciate.

VIT had a highly dynamic localisation within the cell, existing as a single point in extracellular parasites and shortly after invasion before fragmenting as parasite replication was initiated. We also saw limited co-localisation with CPL, a marker of the vacuolar-associated compartment (VAC). The VAC is a dynamic, poorly-defined vesicular compartment which appears to act as a location for ingested protein digestion (Dou et al., 2014; Liu et al., 2014). Localisation of VIT to the VAC fits well with previous observations that VIT is an iron/H^+^ antiporter (Kato et al., 2019; Labarbuta et al., 2017) as the VAC is acidified (Liu et al., 2014; Miranda et al., 2010). In *P. berghei*, VIT-GFP was found to colocalise with markers for the ER and in exogenous expression in *Xenopus*, did not require a proton gradient to transport iron (Slavic et al., 2016). It is possible that this differential localisation was influenced by the tags used (e.g. the presence of GFP can alter the localisation of proteins (Weill et al., 2018), and we appeared to see changes between localisation with HA, Ty and Myc tags). However, the differences in the localisation observed between *T. gondii* and *P. berghei* may also reflect fundamental differences in the biology of these parasites. *Plasmodium* parasites replicate asexually within red blood cells which contain large amounts of iron-containing hemoglobin. For this reason, in *Plasmodium* the vast majority of parasite-associated iron is sequestered as hemozoin crystals within the digestive vacuole (Kapishnikov et al., 2017). In contrast, *T. gondii* lives in the relatively iron-poor environment of nucleated cells. It is this difference in environmental conditions which may be driving the differences in localisation observed. However, the phenotype of VIT in both species suggests a surprising overlap in mechanisms of iron detoxification. Further studies, examining the localisation of VIT in greater detail in *T. gondii* and *Plasmodium* may be helpful to understand the location and dynamics of iron storage in both species.

Our data reveal that VIT levels drop transcriptionally upon iron excess, and at the protein level both upon iron depletion and excess. Transcriptional and post-transcriptional control of iron uptake and storage have been examined in detail in bacteria, yeast, plants and mammalian cells (Andrews et al., 2003; Gao and Dubos, 2021; Li et al., 2008; Philpott et al., 2012; Wang and Pantopoulos, 2011), and was recently also demonstrated in the kinetoplastid parasite *Trypansoma brucei* (Carbajo et al., 2021). However, iron-mediated regulation has not previously been observed in the Apicomplexa. Interestingly, upon excess iron we saw a drop in transcript and protein levels of VIT, the opposite response to the predicted upregulation,as seen in yeast (Li et al., 2008). This suggests that *T. gondii* VIT may have other functions within the cell, however this will be investigated in future work. Recently, expression of an amino acid transporter, ApiAT1, has been shown to be regulated by substrate availability via an upstream small ORF in *T. gondii* (Rajendran et al., 2021). VIT now joins ApiAT1 as the only substrate-regulated transporters identified in the Apicomplexa, although our results suggest that VIT may be regulated at multiple levels, including by modulating localisation. Understanding when and how each method of regulation is used by the parasite, and the pathways controlling this, remains to be unpicked.

It has been shown that in bacteria, yeast, and some plants under stressful conditions, cells will induce iron sparing programs, reducing the production of some iron-containing proteins while prioritising others, often orchestrated at the transcriptional level (Blaby-Haas and Merchant, 2013; Lin et al., 2011; Philpott et al., 2012; Sanvisens et al., 2011). Our findings provide the first evidence of such programs in *T. gondii*. In the absence of VIT, *T. gondii* represses IscU while upregulating IscA. Transcriptional control of IscU by iron levels has been observed in a number of organisms (Garland et al., 1999; Tong and Rouault, 2006) and regulation of an IscA homolog by iron has been observed in bacteria (Andrews et al., 2003). Given the recent interest in iron sulfur biogenesis in *T. gondii* as a target for drug development (Aw et al., 2021; Pamukcu et al., 2021), these findings suggest a currently unknown mechanism regulating this essential metabolic pathway. We also see regulation of RNR, an iron-containing enzyme, and of several genes with predicted DNA or RNA binding motifs. Future studies will examine the response of the parasite to altered iron levels within the cell in more detail to examine potential pathways of iron prioritisation in *T. gondii*.

We found that the increased sensitivity of the ΔVIT parasite to excess iron was almost entirely driven by increased ROS, as ROS levels in the ΔVIT parasites were very high upon treatment with excess iron, and scavenging of ROS by NAC returned parasite iron sensitivity to indistinguishable from the parental line. We believe that under normal conditions, ΔVIT parasites attempted to compensate for the lack of VIT by increasing the activity (although not expression) of catalase and possibly other antioxidants, however this increase was not sufficient to prevent ROS accumulation upon further stresses. Interestingly, loss of CCC1 in yeast is associated with increased oxidative stress (de Llanos et al., 2016; Li et al., 2010) and loss of correct iron storage through ferritin depletion in *Arabidopsis* led to increased oxidative stress, which was partially compensated by increase in antioxidant activity (Ravet et al., 2009; Ruangkiattikul et al., 2012). The importance of iron storage and detoxification across the tree of life highlights the dangers of iron to almost all cells. In *T. gondii*, the response of the parasites to oxidative stress and its role in the lytic life cycle has of been of interest recently (Alves et al., 2021; Augusto et al., 2021) and how iron uptake and storage are linked with these pathways will be an interesting future area of research.

VIT also has an important role in survival in activated macrophages. Activation of macrophages by IFNγ leads to nutrient restriction, including reduced levels of iron (Abreu et al., 2020; Adams et al., 1990; Appelberg, 2006). Further, a previous study on IFNγ-activated enterocytes saw that *T. gondii* growth restriction was reversible by addition of excess iron (Dimier and Bout, 1998). Given that the ΔVIT parasites are more susceptible to acute iron loss, this may be enough to restrict parasite replication, although we cannot rule out that the parasites are more susceptible to the stressful conditions of the phagosome.

In conclusion, we provide the first study of an iron regulated-iron transporter in *T. gondii*. We show that it is not essential for growth under normal conditions, but is vital in detoxification of excess iron. This work provides a basis for the future study of iron within *T. gondii* where many questions remain, including how iron enters the parasite, how it moves between the organelles and the mechanisms of iron-mediated regulation. The conservation of VIT between the Apicomplexa demonstrates that, despite the very different environments that the parasites make home, they utilise some common mechanisms to limit environmental toxicity.

## Materials and Methods

### *T. gondii* and host cell maintenance

*Toxoplasma gondii* tachyzoites were maintained at 37°C with 5% CO_2_ and were grown in human foreskin fibroblasts (HFFs) cultured in Dulbecco’s modified Eagle’s medium (DMEM) that was supplemented with either 3% (D3) or 10% heat-inactivated foetal bovine serum (D10) (FBS), 2mM L-glutamine and 10 μg ml-1 gentamicin.

### *T. gondii* strain construction

The ΔKu80::mNeon and ΔKu80::tdTomato strains were generated by transfection of 40 μg of plasmids containing Cas9 with the required sgRNA (sgRNA sequences are listed in **Table 1**) along with 30 μg of purified PCR product (mNeon green expressing cassette) - amplified using primers p1 and p2 (see **Table 2** for primer sequences) into the RHΔHX line for the ΔKu80::mNeon line and with 30 μg of purified PCR product (tdTomato red expressing cassette) - amplified using primers p3 and p4 for the ΔKu80::tdTomato line. Parasites were transfected into the parental RHΔHX line using a Gene Pulse BioRad electroporator as previously described (Herm-Götz et al., 2007). Fluorescent parasites were isolated by FACS using an Aria III (BD Biosciences), and sorted directly into a 96 well plate pre-seeded with HFFs and incubated at 37°C with 5% CO_2_ for 5-7 days. Positive fluorescent plaques were then identified using microscopy. Absence of *ku80* was confirmed by PCR using primers p7 and p8

**Table 2.**
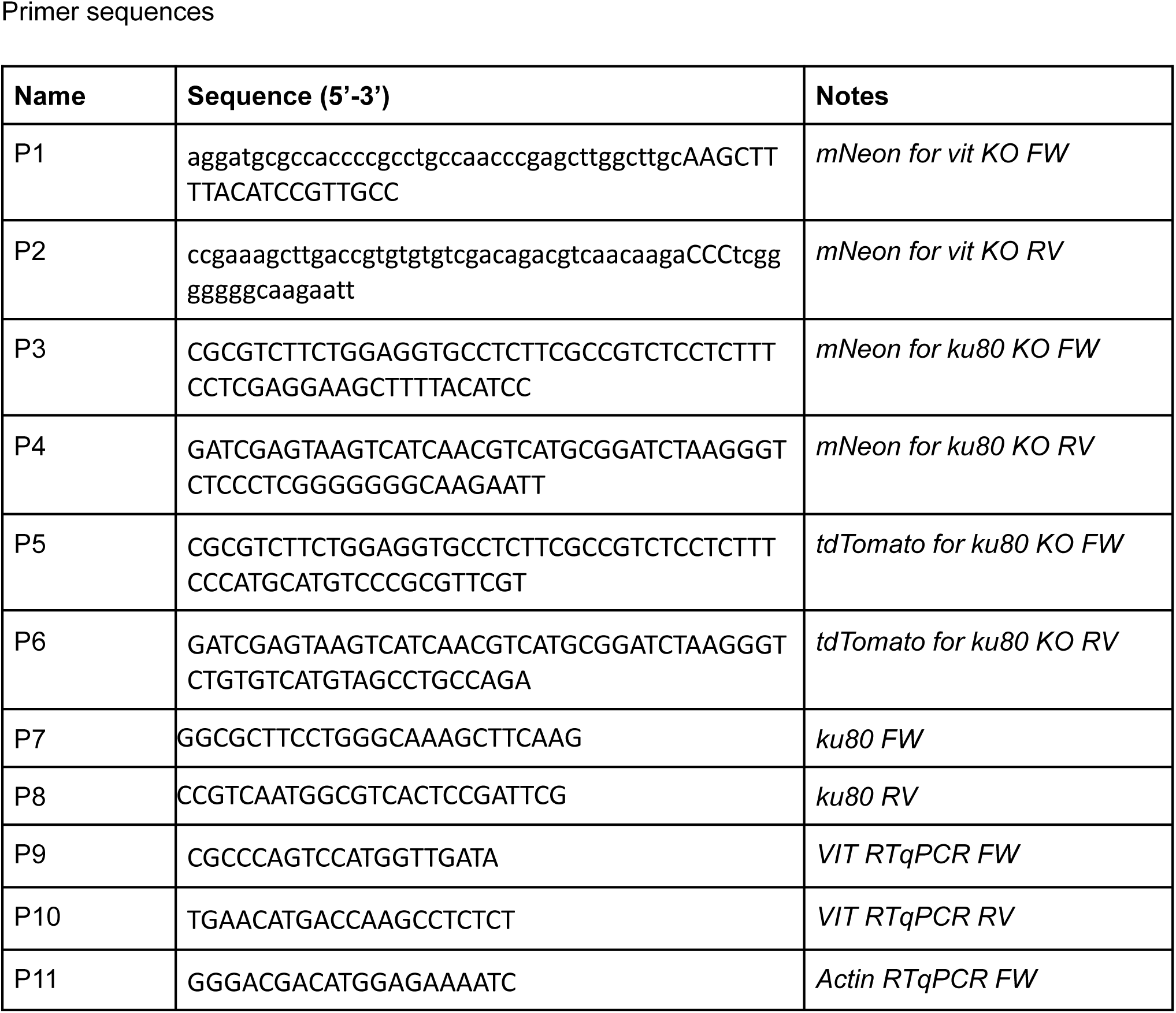
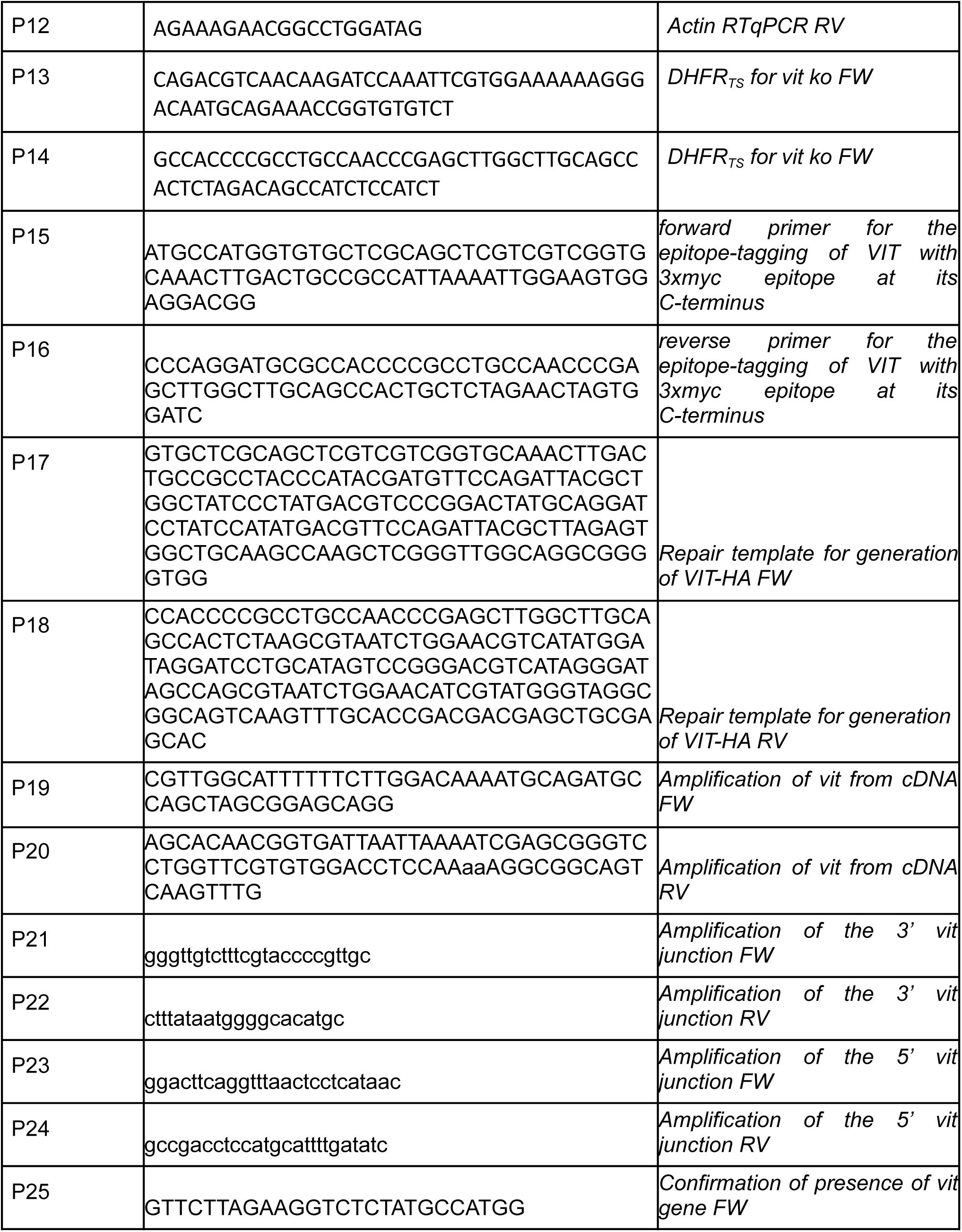
A list of the primer sequences used in this publication

The ΔVIT strain was constructed by co-transfection of two Cas9 guides targeted to the 5’ and 3’ ends of the genomic sequence with a repair template consisting of an mNeonGreen cassette (including a strong SAG1 promotor) in the opposite orientation from *vit*. Parasites were sorted and cloned out as above and integration of the repair cassette confirmed by PCR using primers 21 and 22 for the 3’ junction, primers 23 and 24 for the 5’ junction and 24 and 25 to amplify the endogenous gene The ΔVIT::DHFR_TS_ (Reynolds et al., 2001) was constructed by co-transfection of two Cas9 guides targeted to the 5’ and 3’ ends of the genomic sequence (**Table 1**) with a repair template consisting of the coding region of the pyrimethamine resistance cassette using primers p13 and p14. Transgenic parasites were selected with pyrimethamine, cloned by limiting dilution, and confirmed by Sanger sequencing.

The VIT-HA strain was constructed by transfection of a Cas9 guide targeting the 3’ end of the genomic sequence (**Table 1**) and repaired with an annealed synthetic oligonucleotide (IDT) encoding 40 base pairs of homology and inserting three copies of the HA epitope sequence prior to the stop codon. Fifty million tachyzoites were transfected by electroporation in a 4 mm gap cuvette using a Bio-Rad Gene Pulser II with an exponential decay program set to 1500 V, 25 μF capacitance and no resistance and immediately added to a confluent HFF monolayer in a T25 flask. Parasites were selected with 50 μg/mL phleomycin at 24 h post transfection for 6 h and subsequently added to a confluent HFF monolayer in T25 flask. Parasites were cloned by serial dilution and insertion of the HA tag was confirmed by IFA and western blot.

The VIT-Myc strain was constructed by transfection of a Cas9 guide targeting the 3’ end of the genomic sequence. The 50 bp upstream and downstream homologous regions around the stop codon of the VIT gene were encoded in the forward and reverse primers to generate the repair template by PCR (using premier p15 and p16, **Table 2**), which flanks the homologous regions at both ends of the 3xMyc epitope tag and chloramphenicol resistance cassette. The plasmid encoding the *TgVIT-*targeting guide RNA and Cas9-GFP and the repair template were co-transfected into RHΔ*ku80* parasites. The stop codon of *vit* was replaced by the 3xMyc epitope tag and chloramphenicol resistance cassette. The transfectants were selected through multiple rounds of chloramphenicol to generate a stable population. The VIT-Myc fusion protein was confirmed by immunoblotting analysis.

The VIT-Ty overexpression plasmid was constructed from the MyoAtail-GFP overexpression construct (Harding et al., 2016). The MyoAtail-GFPwas removed by digestion with EcoRI and PacI and *vit* amplified from cDNA using primers 19 and 20 (**Table 2**), which contained a Ty epitope tag. The insert was cloned into the vector using Gibson assembly (NEB) and the correct sequence confirmed by sequencing.

### Immunofluorescence Assays

Indirect immunofluorescence assays (IFA) were performed on both infected HFF monolayers or naturally egressed extracellular tachyzoites. Intracellular *T*. gondii parasites that were grown on HFFs on coverslips were fixed with 4% paraformaldehyde (PFA) for 20 minutes at room temperature and then washed with 1X phosphate buffered saline (PBS). Extracellular parasites for immunofluorescence assays were spun onto poly-L-lysine (Advanced BioMatrix) coated coverslips and fixed with 4% PFA for 20 minutes at room temperature and washed once in PBS. Cells were permeabilized and blocked in PBS/0.2% Triton X-100/2% Bovine Serum Albumin (BSA) (PBS/Triton/BSA), for 30 minutes at room temperature or 4°C overnight. The slides were then incubated in a wet chamber with primary antibodies (rat anti-HA, Merk (11867423001) 1:1000 and guinea pig anti-CDPK, a kind gift of Dr Sebastian Lourido (Whitehead Institute), 1:5000, rabbit anti-CPL, a kind gift from Dr. Vern Carruthers (University of Michigan), 1:500, mouse anti-MYC (LifeTechnologies) 1:400) in the PBS/Triton/BSA mixture for 1 hour at room temperature. After washing, slides were incubated in a wet chamber with the secondary antibodies (Alexa Fluor Goat anti-Rat 594, Invitrogen, 1:1000 and Alexa Fluor Goat anti-Guinea Pig, Invitrogen, 1:1000) in PBS/Triton/BSA for 1 h at room temperature in the dark. After further washes, cells were then mounted onto slides with Fluoromount with DAPI (Southern Biotech). Micrograph images were obtained using a DeltaVision microscope (Applied Precision) and processed and deconvolved using SoftWoRx and FIJI software.

### Plaque assays

Plaque assays were performed as previously described (Sidik et al., 2016). Briefly, 500 parental or ΔVIT parasites were applied onto confluent monolayers of HFF cells. Parasites were allowed to replicate for 7 days undisturbed before being washed with PBS, fixed with ice cold 70 % ethanol and stained using crystal violet stain (12.5 g crystal violet in 125 ml ethanol, diluted in 500 ml 1% ammonium oxalate) for 10 mins at room temperature before washing with distilled H_2_O and imagining.

### Extracellular survival

To assess extracellular survival, parasites were mechanically released from host cells, counted, and diluted to 1 parasite/μl in D3 and incubated at 37 °C for the indicated time. 100 μl of parasites were then placed onto two wells of a 12 well plate of confluent HFF cells and allowed to form plaques (as above) for 5 - 7 days. Plaques were counted and normalised to parasites not incubated extracellularly.

### Competition assay

ΔKu80::mNeon, ΔKu80::tdTomato and ΔVIT extracellular parasites were counted and normalized to 2×10^7^ cells/ml in HBSS + 1% FBS. Equal numbers of ΔKu80::mNeon and ΔKu80::tdTomato parasites, and ΔKu80::tdTomato and ΔVIT parasites were co-inoculated in a 1:1 ratio into host cells with either DMEM alone or DMEM supplemented with 200 μM FAC (2 technical replicates per condition). The mixed populations were subsequently passaged every 2 or 3 days for about 10 days. At every passage, parasites were filtered and collected for analysis using a BD Celesta, data acquired using FACSDiva software (BD Biosciences). Data were processed using FlowJo v10 (BD Biosciences).

### Fluorescent growth assay

Growth assays of fluorescent cells were performed using mNeon-expressing parasites in black, clear bottomed 96 well plates, preseeded with HFFs. Host cells were infected with 1,000 either ΔKu80::mNeon or ΔVIT tachyzoites per well and were allowed to invade for 2 hours at 37 °C with 5% CO_2_. After 2 hours, the media was supplemented with either FAC, DFO, Ars, ZnSO_4_, CuSO_4_, CdCl_2_ at appropriate concentrations and incubated at 37°C with 5% CO_2_ for 4 days. 5 mM N-acetyl-cysteine (NAC, Sigma A7250) was added to all wells where appropriate. Fluorescence (at 594 nm emission) for each well was recorded at 4 days using a PHERAstar FS microplate reader (BMG LabTech). All experiments performed in triplicate wells, at least three independent biological replicates. Uninfected wells served as blanks and fluorescence values were normalised to infected, untreated wells. Dose-response curves and EC_50_s were calculated using GraphPad Prism 9 software.

### ICP-MS

Total parasite-associated iron was quantified using ICP-MS. Confluent HFF monolayers in D150 culture flasks were infected with either mock, parental, or *Δ*VIT::DHFS_TS_ parasites and cultured for 30h. Parasites were harvested by mechanical lysis and filtration through 0.3 µm filters and quantified on a hemocytometer. Parasites were washed with chelexed PBS and digested in 30 % Nitric acid at 85 °C for 3h. Digested samples were diluted in 2% Nitric acid with 0.025% Triton X-100. A four-point calibration curve was generated (5, 10, 25, and 50 ppb Fe) and ^56^Fe was quantified using Scandium (^45^Sc) as an internal standard.

### Alamar blue assay

HFF cells were grown in black opaque 96-well plates until confluent and were treated with indicated concentrations of sodium arsenite (Ars). This was then followed by an Alamar Blue assay with 0.5 mM resazurin at 37 °C with 5% CO_2_ for 4 hours. PHERAstar FS microplate reader (BMG LabTech) was used to determine fluorescence at 530/25 excitation and 590/25 emission. Host cell viability was determined by normalising values to untreated HFFs. The experiment was performed in triplicate for three biological replicates.

### RT-qPCR

RT-qPCR was used to assay relative *vit* expression. Human foreskin fibroblasts were incubated in either D3 or D3 supplemented with ferric ammonium citrate (5 mM) for 6 hours prior to infection with *T. gondii* (RHΔKu80). Parasites were cultured for 24 hours prior to collection by scraping the HFF monolayer, passing the cell suspension through a 26G needle six times and filtration to remove host cell debris. Parasites were then pelleted by centrifugation at 1500 × g for 10 min, the media was then removed and cell pellets frozen and stored at -80 °C. Total RNA was extracted from parasite pellets using the RNAeasy Mini kit (Qiagen) and DNAse I (Invitrogen) treated for 15 mins at room temperature followed by DNAse denaturation at 65 °C for 20 mins. cDNA synthesis was performed with the High Capacity cDNA RT kit (Invitrogen) according to manufacturer’s instructions.

RT-qPCR was carried out on the Applied Biosystems 7500 Real Time PCR system using Power SYBRgreen PCR master mix (Invitrogen), 2 ng of cDNA per reaction and the following cycling conditions: 95 °C for 10 minutes, 40 cycles of 95 °C for 15 seconds, 60 °C for 1 minute. Primer sequences can be found in **Table 2**. Reactions were run in triplicate from three independent biological experiments. Relative fold changes for treated vs. untreated cells was calculated using the 2^-ΔΔ^Ct method (Livak and Schmittgen, 2001) using actin as a housekeeping gene control. Graphpad Prism 9 was used to perform statistical analysis. Specifically, *t* tests testing the null hypothesis - that for each treatment ddCt_average_ = 0, were used to examine the statistical significance of the observed fold changes.

### RNAseq

T75 flasks of human foreskin fibroblasts were incubated in either standard D3, D3 supplemented with 5 mM FAC or 100 μM DFO for 24 h prior to infection. Each T75 was then infected with 6-7 × 10^6^ cells of wild type (RHΔKu80) or ΔVIT. Infected monolayers were cultured at 37 °C, 5% CO_2_ for 24 hours prior to parasite collection. Parasites were pelted by centrifugation at 1500 × g for 10 min and stored at -80 °C as dry pellets until required. RNA was extracted from the pellets using the RNAeasy kit (Qiagen) according to the manufacturer’s instructions. RNA libraries were then prepared using Illumina Stranded mRNA library preparation method and sequenced at 2 × 75 bp to an average of more than 5 million reads per sample. Raw sequencing data (FASTQ format) was processed using the Galaxy public server hosted by EuPathDB (https://veupathdb.globusgenomics.org/). FastQC and Trimmomatic were used for quality control and to remove low quality reads (where Q < 20 across 4bp sliding windows) and adapter sequences (Bolger et al., 2014). The filtered reads were aligned to the *T. gondii* ME49 genome using HITSAT2 (Kim et al., 2015). These sequence alignments were used to identify reads uniquely mapped to annotated genes using HTseq-count. Differential expression analysis was performed in R using DESeq2 (Love et al., 2014). Additional packages EnhancedVolcano (Blighe K et al., 2021), pheatmap and ggplot2 were used to generate plots. All processed data is in Table S2 and raw FASTA files are available at http://www.ncbi.nlm.nih.gov/bioproject/, bioproject ID: PRJNA754376.

### Western blotting

Parasites were filtered and collected by centrifugation at 1500xg for 10 minutes. Samples were lysed with RIPA lysis buffer (150 mM sodium chloride, 1 % Triton X-100, 0.5 % sodium deoxycholate, 0.1 % sodium dodecyl sulfate (SDS) and 50 mM Tris, pH 8.0) for 15 minutes on ice. Samples were then resuspended with 4X SDS loading dye with 5% w/v beta-mercaptoethanol, boiled at 95 °C for 5 minutes and separated on a 10% SDS-PAGE gel for 1.2 h at 160 v. EZ-Run Prestained protein (Fisher) ladder was used as a molecular weight marker. Proteins were then semi-dry transferred to nitrocellulose membrane in Towbin buffer (0.025 M Tris, 0.192 M Glycine, 10 % methanol) for 30 minutes at 190 mA and blocked at room temperature in 5 % milk in 0.1 % Tween/PBS. Blots were then stained with antibodies for 1 h at room temperature: primary rat anti-HA (Merk) at 1:500 and rabbit anti-Tom40 (van Dooren et al., 2016) at 1:2,000 followed by secondary fluorescent antibodies: goat anti-rat coupled to IRDye 800 (1:10,000, LI-COR) and goat anti-rabbit coupled to IRDye 680CW (1:10,000, LI-COR) and visualized using the Odyssey LCX.

### Flow cytometry

CellROX staining was used to quantify ROS as previously described (Matta et al., 2018). Briefly, ΔKu80::mNeonGreen or ΔVIT parasites were untreated or treated with 2 mM FAC overnight, collected and washed once with PBS. Cells were stained with 10 μM CellROX Deep red (Thermo Fisher, C10422) in PBS and incubated in the dark for 1 h. Parasites, including unstrained controls, were then analysed on a BD Celesta analyser and data acquired using FACSDiva software (BD Biosciences). Parasites were gated on forward and side scatter and on green fluorescence. The geometric mean of CellROX fluorescence from at least 5000 parasites on the Cy7 channel was determined using FlowJo v10 software (BD Biosciences) and results from four independent experiments plotted. For MitoNeoD staining, cells were treated and prepared as above and stained with 5 μM for 15 min before analysis, gating as above and reading geometric mean of the Cy5 channel. At least 5000 parasites/sample were analysed and the results represent 4 independent experiments, normalised to the untreated.

### Catalase activity assay

Parental and ΔVIT parasites were untreated or treated overnight with 2 mM FAC and the activity of catalase assessed as described previously (Hadwan, 2018; Portes et al., 2015). Briefly, parasites were collected, filtered, pelted by centrifugation, and resuspended in 80 μl of molecular grade H_2_O with 20 μl of PBS. Parasites were lysed by freeze-thawing three times on dry ice before 50 μl of lysate was mixed with 100 μl of 10 mM H_2_O_2_ and incubated at 37 °C for 2 mins. The lysate was then mixed with 600 μl of cobalt buffer (3.5 mM Co(NO_3_)_2_·6H_2_O, 3.2 mM sodium hexametaphosphate, (NaPO_3_)_6_ and 850 mM sodium bicarbonate) and incubated in the dark at room temperature for 10 mins. Blank samples (only cobalt buffer) and standard (no lysate) controls were included. Absorbance was read at 440 nm using a PHERAstar plate reader and activity was calculated using the following formula:

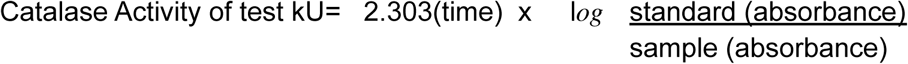

### In gel SOD activity assay

Parasite SOD activity was assessed as previously described with some modifications (Sibley et al., 1986). Parental and ΔVIT parasites were scraped, syringed, filtered, and pelleted. Uninfected host cells were also scraped and syringed but the filtering step was omitted. Cell pellets were resuspended in 100 mM KPO_4_ buffer supplemented with cOmplete, EDTA-free protease inhibitors (Sigma) and incubated on ice for 5 mins. 10 μl was then reserved for western blotting while the remaining was mixed with native loading buffer (2 M sucrose, 0.5% bromophenol blue) and loaded onto a polyacrylamide gel in the abscess of SDS. The gel was ran at 150 v for 2-3 h at 4 °C before being washed gently in water and incubated in development solution (50 ml of 100 mM KPO_4_, 50 mg Nitro Blue Tetrazolium, 7.5 mg riboflavin, 162 μl TEMED) for 45 mins in the dark. The gel was then exposed to light and washed in distilled water 2 - 3 times before imaging.

### Macrophage survival assay

2.5 × 10^6^/ml RAW 264.7 macrophages (ATCC TIB-71) were seeded onto a black, clear-bottomed 96 well plate and allowed to adhere for 2 - 4 h. Macrophages were then activated by the addition of 0.2 μg/ml LPS and 0.1 ng/ml murine recombinant interferon-γ (IFNγ) (both ThermoFisher Scientific) for 24 h. Macrophages were infected with ΔKu80::mNeon or ΔVIT parasites at an MOI of 10 and fluorescence quantified at 48 h post infection using a PHERAstar plate reader. Survival was normalised to the fluorescence of naive macrophages. The experiment was performed in technical triplicate at least three times.

### *In vivo* infection

Groups of five female Swiss Webster (Jackson) female mice, aged 6 weeks, were infected with either 20 or 100 tachyzoites intraperitoneally in 150 µl of PBS. The number of injected parasites was confirmed by plaque assay. Surviving mice were tested for seropositivity by ELISA. Animal studies described here adhere to a protocol approved by the Committee on the Use and Care of Animals of the University of Michigan. Results were plotted in Graphpad Prism 9 and differences in survival assessed by the Log-rank (Mantel-Cox) test.

## Acknowledgments

C.R.H. is funded by a Sir Henry Dale Fellowship from the Wellcome Trust and the Royal Society (213455/Z/18/Z) and a Carnegie Research Incentive Grant (RIG009880) from the Carnegie Trust. D.A and C.R.H are funded by a Lord Kelvin/Adam Smith (LKAS) Fellowship from the University of Glasgow. We acknowledge the University of Michigan Office of the Provost and the University of Michigan Departments of Chemistry and Biophysics for support to A.J.G. We thank Vern Carruthers (University of Michigan), Sebastian Lourido (Whitehead Institute, MIT) and Lilach Sheiner (University of Glasgow) for providing antibodies. The authors gratefully acknowledge the assistance of the Institute of Infection, Immunity and Inflammation Flow Core and Imaging facilities and Glasgow Polyomics for sequencing assistance. We thank Elizabeth Peat for invaluable technical assistance and Yara Aghabi for assistance in image quantification. We thank Tracey Schultz for invaluable assistance in performing the mouse infections.

---

**Supplementary Table 1.** List of all results from RNAseq

## Supplemental figures

**Figure S1.**
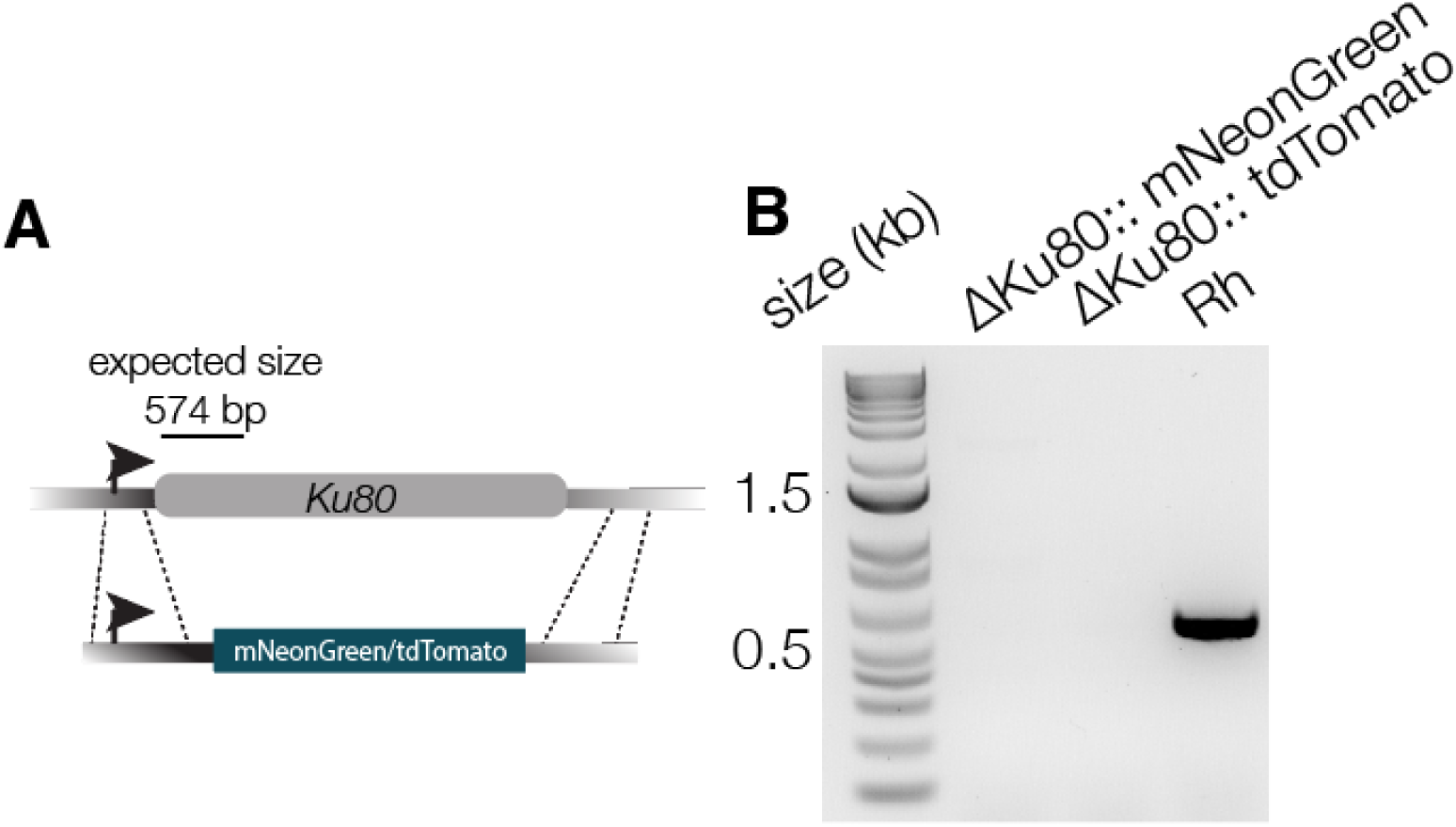
Creation of new ΔKu80 lines expressing mNeonGreen or tdTomato. **A**. Schematic indicating how the endogenous *ku80* gene was replaced with the mNeonGreen or tdTomato cassette. **B**. PCR reactions of region indicated in A, confirming the loss of the endogenous gene.

**Figure S2.**
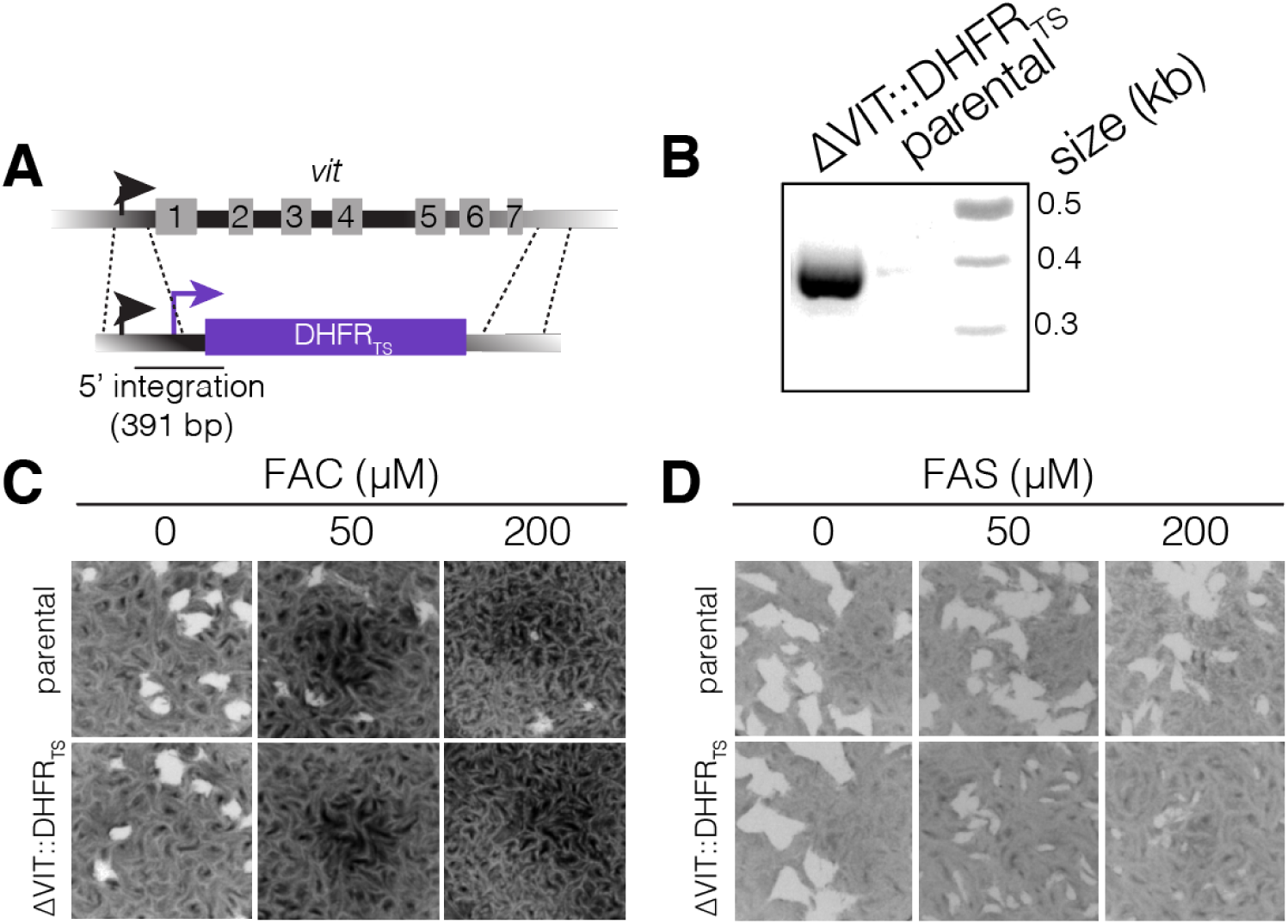
Construction of ΔVIT::DHFR_TS_ parasite line. **A**. Schematic showing how *vit* was replaced with the DHFR cassette. **B**. PCR showing the 5’ integration of the DHFR cassette. Plaque assays showing increased sensitivity of the ΔVIT::DHFR_TS_ line to excess ferric ammonium citrate (FAC) **(C)** and ferrous ammonium sulphate (FAS) **(D)** at the indicated concentration compared to the parental parasite line.

**Figure S3.**
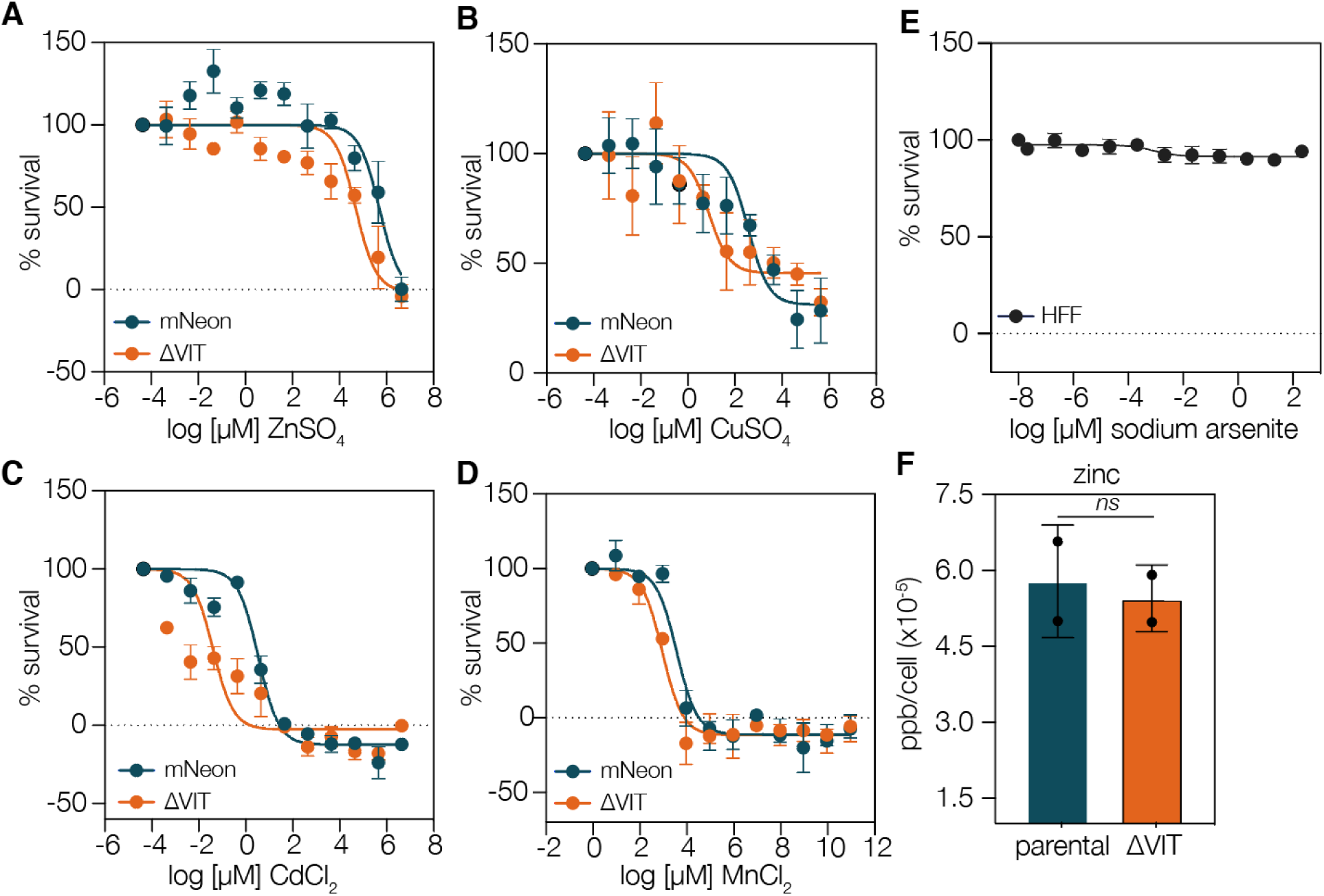
Dose response curves of mNeon and ΔVIT treated with excess metals. As described above, parasites were treated with the indicated concentration of ZnSO_4_ **(A)**, CuSO_4_ **(B)**, CdCl_2_ **(C)**, or MnCl_2_ **(D)**. All experiments are the mean of four (ZnSO4) or three (CuSO_4_, CdCl_2_, and MnCl_2_) independent experiments performed in triplicate, ± SEM. **E**. We assessed survival of uninfected HFF cells at the range of sodium arsenite concentrations used above. No change in fluorescence was seen at the concentrations used. Results are the mean of three independent experiments, ± SEM. **F**. ICP-MS quantifying zinc (66 Zn) from parental and ΔVIT parasites. Each point represents independent biological replicates, performed in technical duplicate. Bars at mean ± SD.

**Figure S4.**
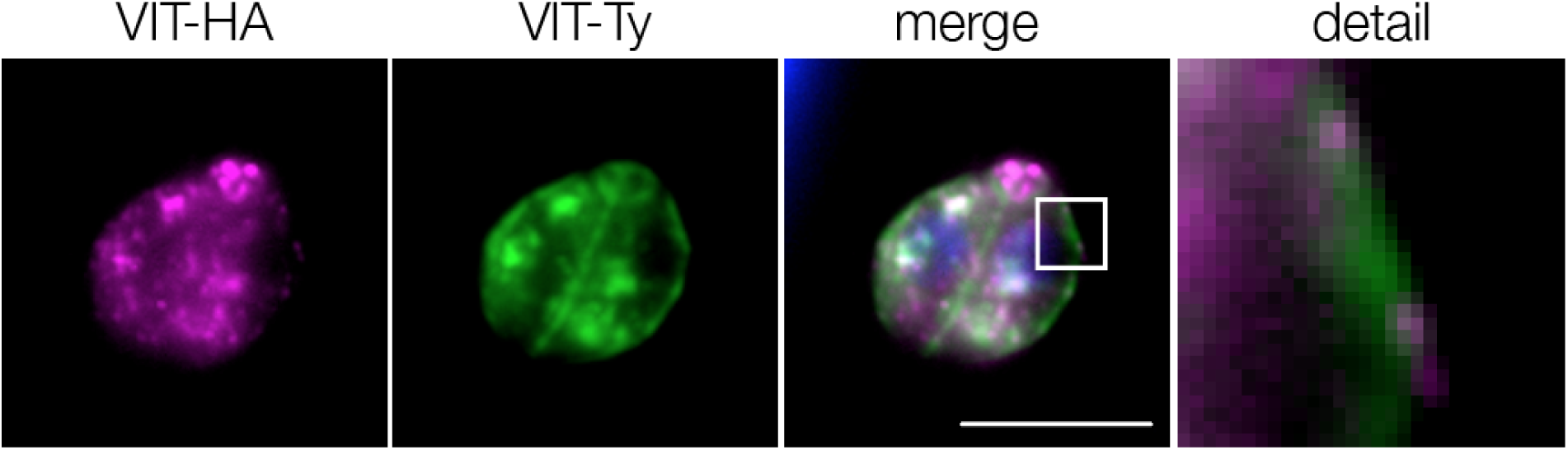
VIT-HA colocalises with VIT-Ty. Colocalisation of endogenously expressed VIT-Ty with overexpressed VIT-HA. Scale bar 5 μm.

**Figure S5.**
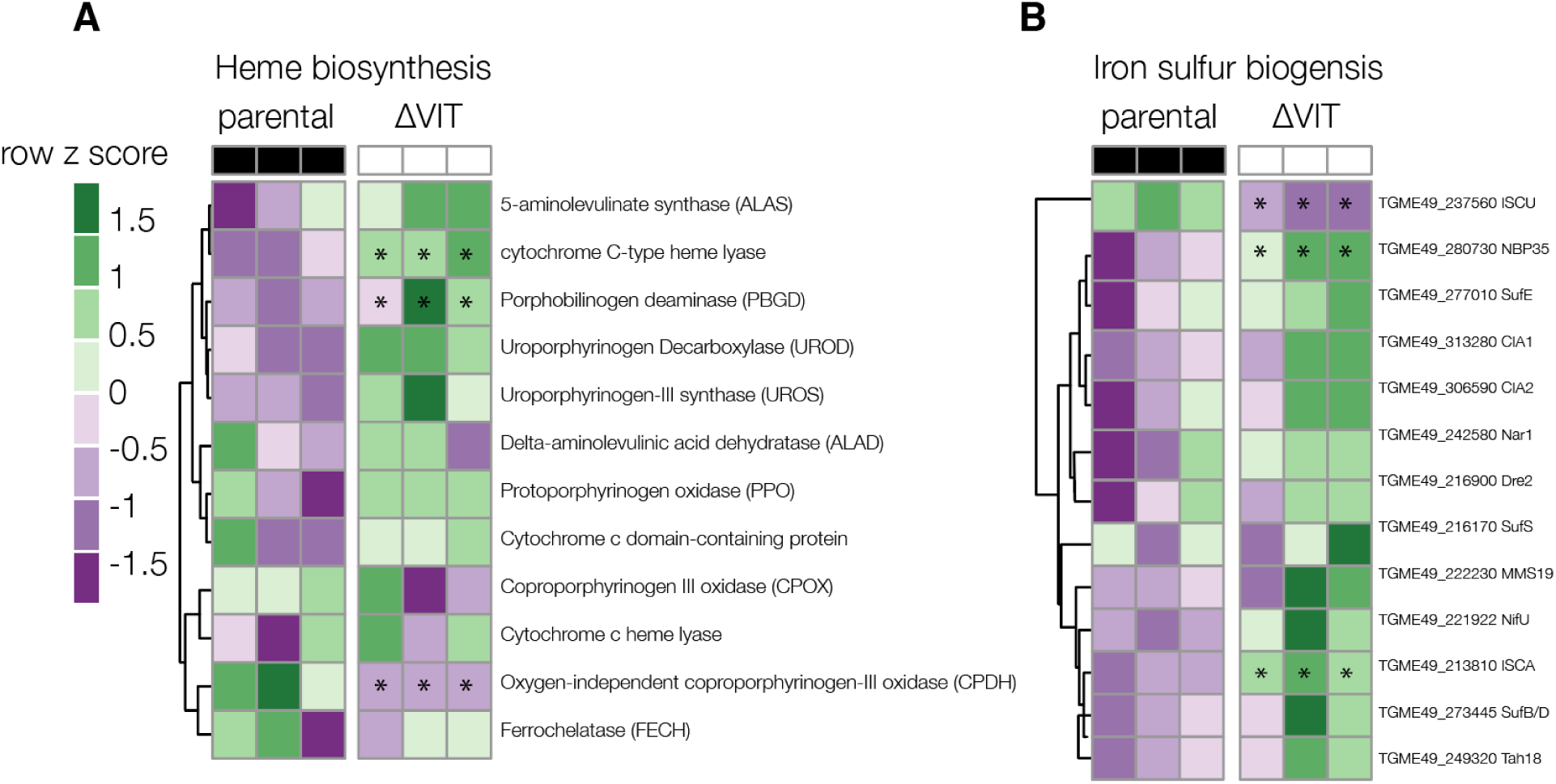
Heatmaps of genes from major iron pathways from parental and ΔVIT parasites. **A**. Genes involved in the heme biosynthesis pathway, read counts normalised across rows. * indicates adj. *p* value of < 0.05 between parental and ΔVIT strain. **B**. Genes involved in Fe-S biogenesis read counts normalised across rows. * indicates adj. *p* value of < 0.05 between parental and ΔVIT strain.

**Figure S6.**
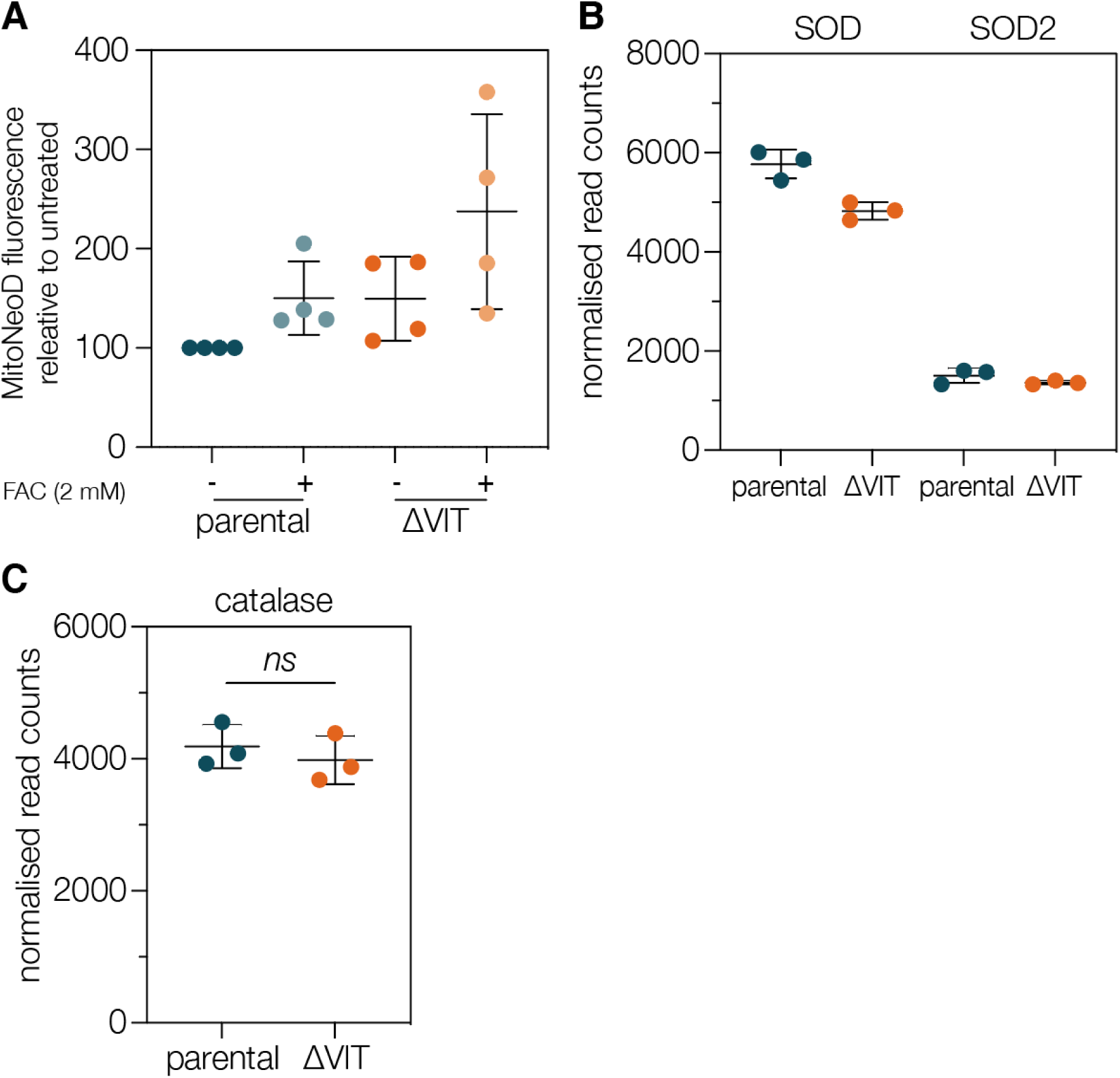
ΔVIT parasites differ in response to oxidative stress. **A**. MitoNeoD fluorescence was used to assess mitochondrial ROS. There was no significant difference under any of the tested conditions, points represent at least 10,000 parasites from four independent experiments, normalised to the parental, untreated line. Bars represent mean ± SD. **B**. Normalised read counts for SOD (TGME49_316310) and SOD2 (TGME49_316330). SOD3 (TGME49_316190) is not expressed under these conditions. Each point represents an independent experiment, performed in duplicate. Line at mean ± SD. **C**. Normalised read counts for catalase (TGME49_232250). Each point represents an independent experiment, performed in duplicate. Line at mean ± SD. *p* value from *t* test.

## Notes

### Competing Interest Statement

The authors have declared no competing interest.

## References

Abreu, R., Essler, L., Giri, P., Quinn, F., 2020. Interferon-gamma promotes iron export in human macrophages to limit intracellular bacterial replication. PLOS ONE 15, e0240949. https://doi.org/10.1371/journal.pone.0240949

Adams, L.B., Hibbs, J.B., Taintor, R.R., Krahenbuhl, J.L., 1990. Microbiostatic effect of murine-activated macrophages for Toxoplasma gondii. Role for synthesis of inorganic nitrogen oxides from L-arginine. J Immunol 144, 2725–2729.

Al-sandaqchi, A.T., Brignell, C., Collingwood, J.F., Geraki, K., Mirkes, E.M., Kong, K., Castellanos, M., May, S.T., Stevenson, C.W., Elsheikha, H.M., 2018. Metallome of cerebrovascular endothelial cells infected with Toxoplasma gondii using μ-XRF imaging and inductively coupled plasma mass spectrometry. Metallomics 10, 1401–1414. https://doi.org/10.1039/C8MT00136G

Alves, E., Benns, H.J., Magnus, L., Dominicus, C., Dobai, T., Blight, J., Wincott, C.J., Child, M.A., 2021. An Extracellular Redox Signal Triggers Calcium Release and Impacts the Asexual Development of Toxoplasma gondii. Front. Cell. Infect. Microbiol. 0. https://doi.org/10.3389/fcimb.2021.728425

Andrews, S.C., Robinson, A.K., Rodríguez-Quiñones, F., 2003. Bacterial iron homeostasis. FEMS Microbiology Reviews 27, 215–237. https://doi.org/10.1016/S0168-6445(03)00055-X

Appelberg, R., 2006. Macrophage nutriprive antimicrobial mechanisms. J Leukoc Biol 79, 1117–1128. https://doi.org/10.1189/jlb.0206079

Arosio, P., Carmona, F., Gozzelino, R., Maccarinelli, F., Poli, M., 2015. The importance of eukaryotic ferritins in iron handling and cytoprotection. Biochem J 472, 1–15. https://doi.org/10.1042/BJ20150787

Arosio, P., Elia, L., Poli, M., 2017. Ferritin, cellular iron storage and regulation. IUBMB Life 69, 414–422. https://doi.org/10.1002/iub.1621

Augusto, L., Martynowicz, J., Amin, P.H., Carlson, K.R., Wek, R.C., Sullivan, W.J., 2021. TgIF2K-B Is an eIF2α Kinase in Toxoplasma gondii That Responds to Oxidative Stress and Optimizes Pathogenicity. mBio 12. https://doi.org/10.1128/mBio.03160-20

Aw, Y.T.V., Seidi, A., Hayward, J.A., Lee, J., Makota, F.V., Rug, M., Dooren, G.G. van, 2021. A key cytosolic iron–sulfur cluster synthesis protein localizes to the mitochondrion of Toxoplasma gondii. Molecular Microbiology 115, 968–985. https://doi.org/10.1111/mmi.14651

Aw, Y.T.V., Seidi, A., Hayward, J.A., Lee, J., Victor Makota, F., Rug, M., van Dooren, G.G., 2020. A key cytosolic iron-sulfur cluster synthesis protein localises to the mitochondrion of Toxoplasma gondii. Mol Microbiol. https://doi.org/10.1111/mmi.14651

Bakouh, N., Bellanca, S., Nyboer, B., Moliner Cubel, S., Karim, Z., Sanchez, C.P., Stein, W.D., Planelles, G., Lanzer, M., 2017. Iron is a substrate of the Plasmodium falciparum chloroquine resistance transporter PfCRT in Xenopus oocytes. J Biol Chem 292, 16109–16121. https://doi.org/10.1074/jbc.M117.805200

Bergmann, A., Floyd, K., Key, M., Dameron, C., Rees, K.C., Thornton, L.B., Whitehead, D.C., Hamza, I., Dou, Z., 2020. Toxoplasma gondii requires its plant-like heme biosynthesis pathway for infection. PLOS Pathogens 16, e1008499. https://doi.org/10.1371/journal.ppat.1008499

Blaby-Haas, C.E., Merchant, S.S., 2013. Iron sparing and recycling in a compartmentalized cell. Current Opinion in Microbiology, Growth and development: eukaryotes/prokaryotes 16, 677–685. https://doi.org/10.1016/j.mib.2013.07.019

Blighe K, Rana S, Lewis M, 2021. EnhancedVolcano: publication-ready volcano plots with enhanced colouring and labeling [WWW Document]. URL https://bioconductor.org/packages/release/bioc/vignettes/EnhancedVolcano/inst/doc/EnhancedVolcano.html#acknowledgments (accessed 8.3.21).

Bolger, A.M., Lohse, M., Usadel, B., 2014. Trimmomatic: a flexible trimmer for Illumina sequence data. Bioinformatics 30, 2114–2120. https://doi.org/10.1093/bioinformatics/btu170

Brembu, T., Jørstad, M., Winge, P., Valle, K.C., Bones, A.M., 2011. Genome-Wide Profiling of Responses to Cadmium in the Diatom Phaeodactylum tricornutum. Environ. Sci. Technol. 45, 7640–7647. https://doi.org/10.1021/es2002259

Carbajo, C.G., Cornell, L.J., Madbouly, Y., Lai, Z., Yates, P.A., Tinti, M., Tiengwe, C., 2021. Novel aspects of iron homeostasis in pathogenic bloodstream form Trypanosoma brucei. PLOS Pathogens 17, e1009696. https://doi.org/10.1371/journal.ppat.1009696

Charvat, R.A., Arrizabalaga, G., 2016. Oxidative stress generated during monensin treatment contributes to altered Toxoplasma gondii mitochondrial function. Scientific Reports 6, 22997. https://doi.org/10.1038/srep22997

Conte, S.S., Walker, E.L., 2011. Transporters contributing to iron trafficking in plants. Mol Plant 4, 464–476. https://doi.org/10.1093/mp/ssr015

de Llanos, R., Martínez-Garay, C.A., Fita-Torró, J., Romero, A.M., Martínez-Pastor, M.T., Puig, S., 2016. Soybean Ferritin Expression in Saccharomyces cerevisiae Modulates Iron Accumulation and Resistance to Elevated Iron Concentrations. Appl. Environ. Microbiol. 82, 3052–3060. https://doi.org/10.1128/AEM.00305-16

Dimier, I.H., Bout, D.T., 1998. Interferon-gamma-activated primary enterocytes inhibit Toxoplasma gondii replication: a role for intracellular iron. Immunology 94, 488–495. https://doi.org/10.1046/j.1365-2567.1998.00553.x

Dou, Z., McGovern, O.L., Cristina, M.D., Carruthers, V.B., 2014. Toxoplasma gondii Ingests and Digests Host Cytosolic Proteins. mBio 5. https://doi.org/10.1128/mBio.01188-14

Epsztejn, S., Glickstein, H., Picard, V., Slotki, I.N., Breuer, W., Beaumont, C., Cabantchik, Z.I., 1999. H-Ferritin Subunit Overexpression in Erythroid Cells Reduces the Oxidative Stress Response and Induces Multidrug Resistance Properties. Blood 94, 3593–3603. https://doi.org/10.1182/blood.V94.10.3593.422k26_3593_3603

Fang, D., Bao, Y., Li, X., Liu, F., Cai, K., Gao, J., Liao, Q., 2010. Effects of iron deprivation on multidrug resistance of leukemic K562 cells. Chemotherapy 56, 9–16. https://doi.org/10.1159/000287352

Florimond, C., Cordonnier, C., Taujale, R., Wel, H. van der, Kannan, N., West, C.M., Blader, I.J., 2019. A Toxoplasma Prolyl Hydroxylase Mediates Oxygen Stress Responses by Regulating Translation Elongation. mBio 10. https://doi.org/10.1128/mBio.00234-19

Galaris, D., Barbouti, A., Pantopoulos, K., 2019. Iron homeostasis and oxidative stress: An intimate relationship. Biochim Biophys Acta Mol Cell Res 1866, 118535. https://doi.org/10.1016/j.bbamcr.2019.118535

Gao, F., Dubos, C., 2021. Transcriptional integration of plant responses to iron availability. Journal of Experimental Botany 72, 2056–2070. https://doi.org/10.1093/jxb/eraa556

Garland, S.A., Hoff, K., Vickery, L.E., Culotta, V.C., 1999. Saccharomyces cerevisiae ISU1 and ISU2: members of a well-conserved gene family for iron-sulfur cluster assembly Edited by S. Reed. Journal of Molecular Biology 294, 897–907. https://doi.org/10.1006/jmbi.1999.3294

Gollhofer, J., Timofeev, R., Lan, P., Schmidt, W., Buckhout, T.J., 2014. Vacuolar-Iron-Transporter1-Like Proteins Mediate Iron Homeostasis in Arabidopsis. PLOS ONE 9, e110468. https://doi.org/10.1371/journal.pone.0110468

Hadwan, M.H., 2018. Simple spectrophotometric assay for measuring catalase activity in biological tissues. BMC Biochemistry 19, 7. https://doi.org/10.1186/s12858-018-0097-5

Harding, C.R., Egarter, S., Gow, M., Jiménez-Ruiz, E., Ferguson, D.J.P., Meissner, M., 2016. Gliding Associated Proteins Play Essential Roles during the Formation of the Inner Membrane Complex of Toxoplasma gondii. PLoS Pathog. 12, e1005403. https://doi.org/10.1371/journal.ppat.1005403

Herm-Götz, A., Agop-Nersesian, C., Münter, S., Grimley, J.S., Wandless, T.J., Frischknecht, F., Meissner, M., 2007. Rapid control of protein level in the apicomplexan Toxoplasma gondii. Nat Methods 4, 1003–1005. https://doi.org/10.1038/nmeth1134

Hu, Y., Li, J., Lou, B., Wu, R., Wang, G., Lu, C., Wang, H., Pi, J., Xu, Y., 2020. The Role of Reactive Oxygen Species in Arsenic Toxicity. Biomolecules 10. https://doi.org/10.3390/biom10020240

Imlay, J.A., Chin, S.M., Linn, S., 1988. Toxic DNA damage by hydrogen peroxide through the Fenton reaction in vivo and in vitro. Science 240, 640–642. https://doi.org/10.1126/science.2834821

Jasinski, M., Sudre, D., Schansker, G., Schellenberg, M., Constant, S., Martinoia, E., Bovet, L., 2008. AtOSA1, a Member of the Abc1-like Family, as a New Factor in Cadmium and Oxidative Stress Response. Plant Physiology 147, 719–731.

Kapishnikov, S., Grolimund, D., Schneider, G., Pereiro, E., McNally, J.G., Als-Nielsen, J., Leiserowitz, L., 2017. Unraveling heme detoxification in the malaria parasite by in situ correlative X-ray fluorescence microscopy and soft X-ray tomography. Scientific Reports 7, 7610. https://doi.org/10.1038/s41598-017-06650-w

Kato, T., Kumazaki, K., Wada, M., Taniguchi, R., Nakane, T., Yamashita, K., Hirata, K., Ishitani, R., Ito, K., Nishizawa, T., Nureki, O., 2019. Crystal structure of plant vacuolar iron transporter VIT1. Nature Plants 5, 308–315. https://doi.org/10.1038/s41477-019-0367-2

Kim, D., Langmead, B., Salzberg, S.L., 2015. HISAT: a fast spliced aligner with low memory requirements. Nat Methods 12, 357–360. https://doi.org/10.1038/nmeth.3317

Kim, S.A., Punshon, T., Lanzirotti, A., Li, L., Alonso, J.M., Ecker, J.R., Kaplan, J., Guerinot, M.L., 2006. Localization of Iron in Arabidopsis Seed Requires the Vacuolar Membrane Transporter VIT1. Science 314, 1295–1298. https://doi.org/10.1126/science.1132563

Kloehn, J., Harding, C.R., Soldati-Favre, D., 2020. Supply and demand—heme synthesis, salvage and utilization by Apicomplexa. The FEBS Journal n/a. https://doi.org/10.1111/febs.15445

Koenderink, J.B., Kavishe, R.A., Rijpma, S.R., Russel, F.G.M., 2010. The ABCs of multidrug resistance in malaria. Trends Parasitol 26, 440–446. https://doi.org/10.1016/j.pt.2010.05.002

Krishnan, A., Kloehn, J., Lunghi, M., Chiappino-Pepe, A., Waldman, B.S., Nicolas, D., Varesio, E., Hehl, A., Lourido, S., Hatzimanikatis, V., Soldati-Favre, D., 2020. Functional and Computational Genomics Reveal Unprecedented Flexibility in Stage-Specific Toxoplasma Metabolism. Cell Host & Microbe 27, 290-306.e11. https://doi.org/10.1016/j.chom.2020.01.002

Kwok, L.Y., Schlüter, D., Clayton, C., Soldati, D., 2004. The antioxidant systems in Toxoplasma gondii and the role of cytosolic catalase in defence against oxidative injury. Molecular Microbiology 51, 47–61. https://doi.org/10.1046/j.1365-2958.2003.03823.x

Labarbuta, P., Duckett, K., Botting, C.H., Chahrour, O., Malone, J., Dalton, J.P., Law, C.J., 2017. Recombinant vacuolar iron transporter family homologue PfVIT from human malaria-causing Plasmodium falciparum is a Fe 2+ /H + exchanger. Scientific Reports 7, 42850. https://doi.org/10.1038/srep42850

Lee, W., Ha, J.-M., Sugiyama, Y., 2020. Post-translational regulation of the major drug transporters in the families of organic anion transporters and organic anion-transporting polypeptides. J Biol Chem 295, 17349–17364. https://doi.org/10.1074/jbc.REV120.009132

Li, L., Bagley, D., Ward, D.M., Kaplan, J., 2008. Yap5 Is an Iron-Responsive Transcriptional Activator That Regulates Vacuolar Iron Storage in Yeast. Mol Cell Biol 28, 1326–1337. https://doi.org/10.1128/MCB.01219-07

Li, L., Chen, O.S., McVey Ward, D., Kaplan, J., 2001. CCC1 is a transporter that mediates vacuolar iron storage in yeast. J Biol Chem 276, 29515–29519. https://doi.org/10.1074/jbc.M103944200

Li, L., Murdock, G., Bagley, D., Jia, X., Ward, D.M., Kaplan, J., 2010. Genetic Dissection of a Mitochondria-Vacuole Signaling Pathway in Yeast Reveals a Link between Chronic Oxidative Stress and Vacuolar Iron Transport. J. Biol. Chem. 285, 10232–10242. https://doi.org/10.1074/jbc.M109.096859

Lin, H., Li, L., Jia, X., Ward, D.M., Kaplan, J., 2011. Genetic and biochemical analysis of high iron toxicity in yeast: iron toxicity is due to the accumulation of cytosolic iron and occurs under both aerobic and anaerobic conditions. J Biol Chem 286, 3851–3862. https://doi.org/10.1074/jbc.M110.190959

Liu, J., Pace, D., Dou, Z., King, T.P., Guidot, D., Li, Z.-H., Carruthers, V.B., Moreno, S.N.J., 2014. A Vacuolar-H+-Pyrophosphatase (TgVP1) is Required for Microneme Secretion, Host Cell Invasion, and Extracellular Survival of Toxoplasma gondii. Mol Microbiol 93, 698–712. https://doi.org/10.1111/mmi.12685

Livak, K.J., Schmittgen, T.D., 2001. Analysis of relative gene expression data using real-time quantitative PCR and the 2(-Delta Delta C(T)) Method. Methods 25, 402–408. https://doi.org/10.1006/meth.2001.1262

Love, M.I., Huber, W., Anders, S., 2014. Moderated estimation of fold change and dispersion for RNA-seq data with DESeq2. Genome Biology 15, 550. https://doi.org/10.1186/s13059-014-0550-8

Lykens, J.E., Terrell, C.E., Zoller, E.E., Divanovic, S., Trompette, A., Karp, C.L., Aliberti, J., Flick, M.J., Jordan, M.B., 2010. Mice with a selective impairment of IFN-gamma signaling in macrophage lineage cells demonstrate the critical role of IFN-gamma-activated macrophages for the control of protozoan parasitic infections in vivo. J Immunol 184, 877–885. https://doi.org/10.4049/jimmunol.0902346

Matta, S.K., Patten, K., Wang, Q., Kim, B.-H., MacMicking, J.D., Sibley, L.D., 2018. NADPH Oxidase and Guanylate Binding Protein 5 Restrict Survival of Avirulent Type III Strains of Toxoplasma gondii in Naive Macrophages. mBio 9, e01393–18. https://doi.org/10.1128/mBio.01393-18

Miranda, K., Pace, D.A., Cintron, R., Rodrigues, J.C.F., Fang, J., Smith, A., Rohloff, P., Coelho, E., de Haas, F., de Souza, W., Coppens, I., Sibley, L.D., Moreno, S.N.J., 2010. Characterization of a novel organelle in Toxoplasma gondii with similar composition and function to the plant vacuole. Mol. Microbiol. 76, 1358–1375. https://doi.org/10.1111/j.1365-2958.2010.07165.x

Odberg-Ferragut, C., Renault, J.P., Viscogliosi, E., Toursel, C., Briche, I., Engels, A., Lepage, G., Morgenstern-Badarau, I., Camus, D., Tomavo, S., Dive, D., 2000. Molecular cloning, expression analysis and iron metal cofactor characterisation of a superoxide dismutase from Toxoplasma gondii. Mol Biochem Parasitol 106, 121–129. https://doi.org/10.1016/s0166-6851(99)00211-x

Pamukcu, S., Cerutti, A., Hem, S., Rofidal, V., Besteiro, S., 2021. Differential contribution of two organelles of endosymbiotic origin to iron-sulfur cluster synthesis in Toxoplasma. bioRxiv 2021.01.28.428257. https://doi.org/10.1101/2021.01.28.428257

Philpott, C.C., Leidgens, S., Frey, A.G., 2012. Metabolic remodeling in iron-deficient fungi. Biochim Biophys Acta 1823, 1509–1520. https://doi.org/10.1016/j.bbamcr.2012.01.012

Portes, J.A., Souza, T.G., Santos, T.A.T. dos Silva, L.L.R., da Ribeiro, T.P., Pereira, M.D., Horn, A., Fernandes, C., DaMatta, R.A., Souza, W., de Seabra, S.H., 2015. Reduction of Toxoplasma gondii Development Due to Inhibition of Parasite Antioxidant Enzymes by a Dinuclear Iron(III) Compound. Antimicrobial Agents and Chemotherapy 59, 7374–7386. https://doi.org/10.1128/AAC.00057-15

Rajendran, E., Clark, M., Goulart, C., Steinhöfel, B., Tjhin, E.T., Gross, S., Smith, N.C., Kirk, K., Dooren, G.G. van, 2021. Substrate-mediated regulation of the arginine transporter of Toxoplasma gondii. PLOS Pathogens 17, e1009816. https://doi.org/10.1371/journal.ppat.1009816

Ravet, K., Touraine, B., Boucherez, J., Briat, J.-F., Gaymard, F., Cellier, F., 2009. Ferritins control interaction between iron homeostasis and oxidative stress in Arabidopsis. The Plant Journal 57, 400–412. https://doi.org/10.1111/j.1365-313X.2008.03698.x

Reynolds, M.G., Oh, J., Roos, D.S., 2001. In Vitro Generation of Novel Pyrimethamine Resistance Mutations in the Toxoplasma gondii Dihydrofolate Reductase. Antimicrob Agents Chemother 45, 1271–1277. https://doi.org/10.1128/AAC.45.4.1271-1277.2001

Ruangkiattikul, N., Bhubhanil, S., Chamsing, J., Niamyim, P., Sukchawalit, R., Mongkolsuk, S., 2012. Agrobacterium tumefaciens membrane-bound ferritin plays a role in protection against hydrogen peroxide toxicity and is negatively regulated by the iron response regulator. FEMS Microbiology Letters 329, 87–92. https://doi.org/10.1111/j.1574-6968.2012.02509.x

Sanvisens, N., Bañó, M.C., Huang, M., Puig, S., 2011. Regulation of ribonucleotide reductase in response to iron deficiency. Mol Cell 44, 759–769. https://doi.org/10.1016/j.molcel.2011.09.021

Sehgal, R., Goyal, K., Sehgal, A., 2012. Trichomoniasis and lactoferrin: future prospects. Infect Dis Obstet Gynecol 2012, 536037. https://doi.org/10.1155/2012/536037

Shafik, S.H., Cobbold, S.A., Barkat, K., Richards, S.N., Lancaster, N.S., Llinás, M., Hogg, S.J., Summers, R.L., McConville, M.J., Martin, R.E., 2020. The natural function of the malaria parasite’s chloroquine resistance transporter. Nature Communications 11, 3922. https://doi.org/10.1038/s41467-020-17781-6

Shaikh, Z.A., Vu, T.T., Zaman, K., 1999. Oxidative Stress as a Mechanism of Chronic Cadmium-Induced Hepatotoxicity and Renal Toxicity and Protection by Antioxidants. Toxicology and Applied Pharmacology 154, 256–263. https://doi.org/10.1006/taap.1998.8586

Sharma, P., Tóth, V., Hyland, E.M., Law, C.J., 2021. Characterization of the substrate binding site of an iron detoxifying membrane transporter from Plasmodium falciparum. Malar J 20, 295. https://doi.org/10.1186/s12936-021-03827-7

Shchepinova, M.M., Cairns, A.G., Prime, T.A., Logan, A., James, A.M., Hall, A.R., Vidoni, S., Arndt, S., Caldwell, S.T., Prag, H.A., Pell, V.R., Krieg, T., Mulvey, J.F., Yadav, P., Cobley, J.N., Bright, T.P., Senn, H.M., Anderson, R.F., Murphy, M.P., Hartley, R.C., 2017. MitoNeoD: A Mitochondria-Targeted Superoxide Probe. Cell Chem Biol 24, 1285-1298.e12. https://doi.org/10.1016/j.chembiol.2017.08.003

Sibley, L.D., Lawson, R., Weidner, E., 1986. Superoxide dismutase and catalase in Toxoplasma gondii. Mol Biochem Parasitol 19, 83–87. https://doi.org/10.1016/0166-6851(86)90069-1

Sidik, S.M., Huet, D., Ganesan, S.M., Huynh, M.-H., Wang, T., Nasamu, A.S., Thiru, P., Saeij, J.P.J., Carruthers, V.B., Niles, J.C., Lourido, S., 2016. A Genome-wide CRISPR Screen in Toxoplasma Identifies Essential Apicomplexan Genes. Cell 166, 1423-1435.e12. https://doi.org/10.1016/j.cell.2016.08.019

Slavic, K., Krishna, S., Lahree, A., Bouyer, G., Hanson, K.K., Vera, I., Pittman, J.K., Staines, H.M., Mota, M.M., 2016. A vacuolar iron-transporter homologue acts as a detoxifier in Plasmodium. Nature Communications 7, 10403. https://doi.org/10.1038/ncomms10403

Söding, J., Biegert, A., Lupas, A.N., 2005. The HHpred interactive server for protein homology detection and structure prediction. Nucleic Acids Res 33, W244–W248. https://doi.org/10.1093/nar/gki408

Sorribes-Dauden, R., Peris, D., Martínez-Pastor, M.T., Puig, S., 2020. Structure and function of the vacuolar Ccc1/VIT1 family of iron transporters and its regulation in fungi. Computational and Structural Biotechnology Journal 18, 3712–3722. https://doi.org/10.1016/j.csbj.2020.10.044

Templeton, D.M., Liu, Y., 2010. Multiple roles of cadmium in cell death and survival. Chem Biol Interact 188, 267–275. https://doi.org/10.1016/j.cbi.2010.03.040

Thornton, L.B., Teehan, P., Floyd, K., Cochrane, C., Bergmann, A., Riegel, B., Stasic, A.J., Cristina, M.D., Moreno, S.N.J., Roepe, P.D., Dou, Z., 2019. An ortholog of Plasmodium falciparum chloroquine resistance transporter (PfCRT) plays a key role in maintaining the integrity of the endolysosomal system in Toxoplasma gondii to facilitate host invasion. PLOS Pathogens 15, e1007775. https://doi.org/10.1371/journal.ppat.1007775

Tong, W.-H., Rouault, T.A., 2006. Functions of mitochondrial ISCU and cytosolic ISCU in mammalian iron-sulfur cluster biogenesis and iron homeostasis. Cell Metabolism 3, 199–210. https://doi.org/10.1016/j.cmet.2006.02.003

van Dooren, G.G., Yeoh, L.M., Striepen, B., McFadden, G.I., 2016. The Import of Proteins into the Mitochondrion of Toxoplasma gondii. J. Biol. Chem. 291, 19335–19350. https://doi.org/10.1074/jbc.M116.725069

Wang, J., Pantopoulos, K., 2011. Regulation of cellular iron metabolism. Biochem J 434, 365–381. https://doi.org/10.1042/BJ20101825

Wang, Y., Sangaré, L.O., Paredes-Santos, T.C., Hassan, M.A., Krishnamurthy, S., Furuta, A.M., Markus, B.M., Lourido, S., Saeij, J.P.J., 2020. Genome-wide screens identify Toxoplasma gondii determinants of parasite fitness in IFNγ-activated murine macrophages. Nat Commun 11, 5258. https://doi.org/10.1038/s41467-020-18991-8

Warring, S.D., Dou, Z., Carruthers, V.B., McFadden, G.I., van Dooren, G.G., 2014. Characterization of the Chloroquine Resistance Transporter Homologue in Toxoplasma gondii. Eukaryot Cell 13, 1360–1370. https://doi.org/10.1128/EC.00027-14

Weill, U., Yofe, I., Sass, E., Stynen, B., Davidi, D., Natarajan, J., Ben-Menachem, R., Avihou, Z., Goldman, O., Harpaz, N., Chuartzman, S., Kniazev, K., Knoblach, B., Laborenz, J., Boos, F., Kowarzyk, J., Ben-Dor, S., Zalckvar, E., Herrmann, J.M., Rachubinski, R.A., Pines, O., Rapaport, D., Michnick, S.W., Levy, E.D., Schuldiner, M., 2018. Genome-wide SWAp-Tag yeast libraries for proteome exploration. Nat Methods 15, 617–622. https://doi.org/10.1038/s41592-018-0044-9

Xue, J., Jiang, W., Chen, Y., Gong, F., Wang, M., Zeng, P., Xia, C., Wang, Q., Huang, K., 2017. Thioredoxin reductase from Toxoplasma gondii: an essential virulence effector with antioxidant function. The FASEB Journal 31, 4447–4457. https://doi.org/10.1096/fj.201700008R

Zaidi, A., Singh, K.P., Ali, V., 2017. Leishmania and its quest for iron: An update and overview. Molecular and Biochemical Parasitology 211, 15–25. https://doi.org/10.1016/j.molbiopara.2016.12.004

Zhang, Y., Xu, Y.-H., Yi, H.-Y., Gong, J.-M., 2012. Vacuolar membrane transporters OsVIT1 and OsVIT2 modulate iron translocation between flag leaves and seeds in rice. Plant J 72, 400–410. https://doi.org/10.1111/j.1365-313X.2012.05088.x

